# Variants in tubule epithelial regulatory elements mediate most heritable differences in human kidney function

**DOI:** 10.1101/2024.06.18.599625

**Authors:** Gabriel B. Loeb, Pooja Kathail, Richard Shuai, Ryan Chung, Reinier J. Grona, Sailaja Peddada, Volkan Sevim, Scot Federman, Karl Mader, Audrey Chu, Jonathan Davitte, Juan Du, Alexander R. Gupta, Chun Jimmie Ye, Shawn Shafer, Laralynne Przybyla, Radu Rapiteanu, Nilah Ioannidis, Jeremy F. Reiter

## Abstract

Kidney disease is highly heritable; however, the causal genetic variants, the cell types in which these variants function, and the molecular mechanisms underlying kidney disease remain largely unknown. To identify genetic loci affecting kidney function, we performed a GWAS using multiple kidney function biomarkers and identified 462 loci. To begin to investigate how these loci affect kidney function, we generated single-cell chromatin accessibility (scATAC-seq) maps of the human kidney and identified candidate *cis*-regulatory elements (cCREs) for kidney podocytes, tubule epithelial cells, and kidney endothelial, stromal, and immune cells. Kidney tubule epithelial cCREs explained 58% of kidney function SNP-heritability and kidney podocyte cCREs explained an additional 6.5% of SNP-heritability. In contrast, little kidney function heritability was explained by kidney endothelial, stromal, or immune cell-specific cCREs. Through functionally informed fine-mapping, we identified putative causal kidney function variants and their corresponding cCREs. Using kidney scATAC-seq data, we created a deep learning model (which we named ChromKid) to predict kidney cell type-specific chromatin accessibility from sequence. ChromKid and allele specific kidney scATAC-seq revealed that many fine-mapped kidney function variants locally change chromatin accessibility in tubule epithelial cells. Enhancer assays confirmed that fine-mapped kidney function variants alter tubule epithelial regulatory element function. To map the genes which these regulatory elements control, we used CRISPR interference (CRISPRi) to target these regulatory elements in tubule epithelial cells and assessed changes in gene expression. CRISPRi of enhancers harboring kidney function variants regulated *NDRG1* and *RBPMS* expression. Thus, inherited differences in tubule epithelial *NDRG1* and *RBPMS* expression may predispose to kidney disease in humans. We conclude that genetic variants affecting tubule epithelial regulatory element function account for most SNP-heritability of human kidney function. This work provides an experimental approach to identify the variants, regulatory elements, and genes involved in polygenic disease.

## INTRODUCTION

Kidney function—the capacity of the kidneys to filter the blood—maintains water, solute, and metabolic waste homeostasis in vertebrates^1^. Chronic kidney disease, defined by persistent decreased kidney function, affects 9% of humans, is the twelfth leading global cause of death, and accounts for an increasing fraction deaths^2^. Lack of understanding of the mechanisms that contribute to kidney disease have hindered development of effective therapies.

Kidney disease and function is highly heritable^3–5^. Genome-wide association studies (GWAS) have identified hundreds of loci associated with biomarkers of kidney function^6–9^. However, for most loci, the causal variants, affected genes, and cells through which the genetic effects are manifested remain unknown.

Clinically, kidney function is estimated in humans using serum biomarkers, substances that are cleared by the kidney and accumulate in the blood when kidney function declines^10^. Genetic variants associated with serum creatinine levels (the most commonly used biomarker of kidney function) are enriched in kidney cCREs^8,11^ with recent work highlighting specific enrichment within kidney proximal tubule cCREs^12,13^. Because individual biomarkers like serum creatinine levels reflect both kidney function and creatinine metabolism, incorporating multiple biomarkers provide more accurate estimates of kidney function^14,15^.

In this study, we systematically identify the cell types, variants, regulatory elements, and genes involved in kidney function. We conducted a GWAS using two serum biomarkers to more accurately estimate kidney function. We identified cCRES specific to different kidney cell types and used these data for functionally-informed variant fine-mapping and development of a sequence-to-accessibility machine learning model, ChromKid. Fine-mapping informed by cell type-specific chromatin accessibility identified variants likely to affect kidney function. Interrogating how cell type-specific cCREs contribute to kidney function heritability indicated the relative contributions of multiple cell types to kidney function. Kidney function heritability was predominantly determined by variants in proximal tubule, distal tubule and podocyte cCREs. A combination of experiment and ChromKid-based prediction revealed that many of these variants affect tubule epithelial chromatin accessibility and regulatory element function. Targeting of regulatory elements containing fine-mapped kidney function variants using CRISPRi revealed regulation of genes not previously implicated in human kidney function. Thus, identification of trait-relevant cell type-specific cCREs, fine-mapping, machine learning, and regulatory element silencing revealed variants, cell types and genes underlying inherited differences in human kidney function.

## RESULTS

### Kidney function heritability is enriched in kidney candidate *cis*-regulatory elements

Kidney function is measured using the glomerular filtration rate (GFR), which is estimated from the concentration of molecules cleared by the kidneys. Previous GWAS have been performed using a single biomarker to calculate estimated GFR (eGFR)^6,7^. eGFR based on both creatinine (cr) and cystatin C (cys) levels (eGFR_cr-cys_) more accurately reflects kidney function^14,15^. Therefore, we performed a GWAS of eGFR_cr-cys_ in individuals of European ancestry within the UK Biobank. This GWAS identified 462 autosomal loci (Figure 1a, Supplementary Table 1). 27 of these loci were not identified through estimates of kidney function using either single biomarker. These 27 loci included *FOXP1*, *RBPJ*, *APOC3*, *IRS2* and *BDKRB2,* genes previously shown to affect kidney injury or development in humans or model organisms^16–21^.

**Figure 1:**
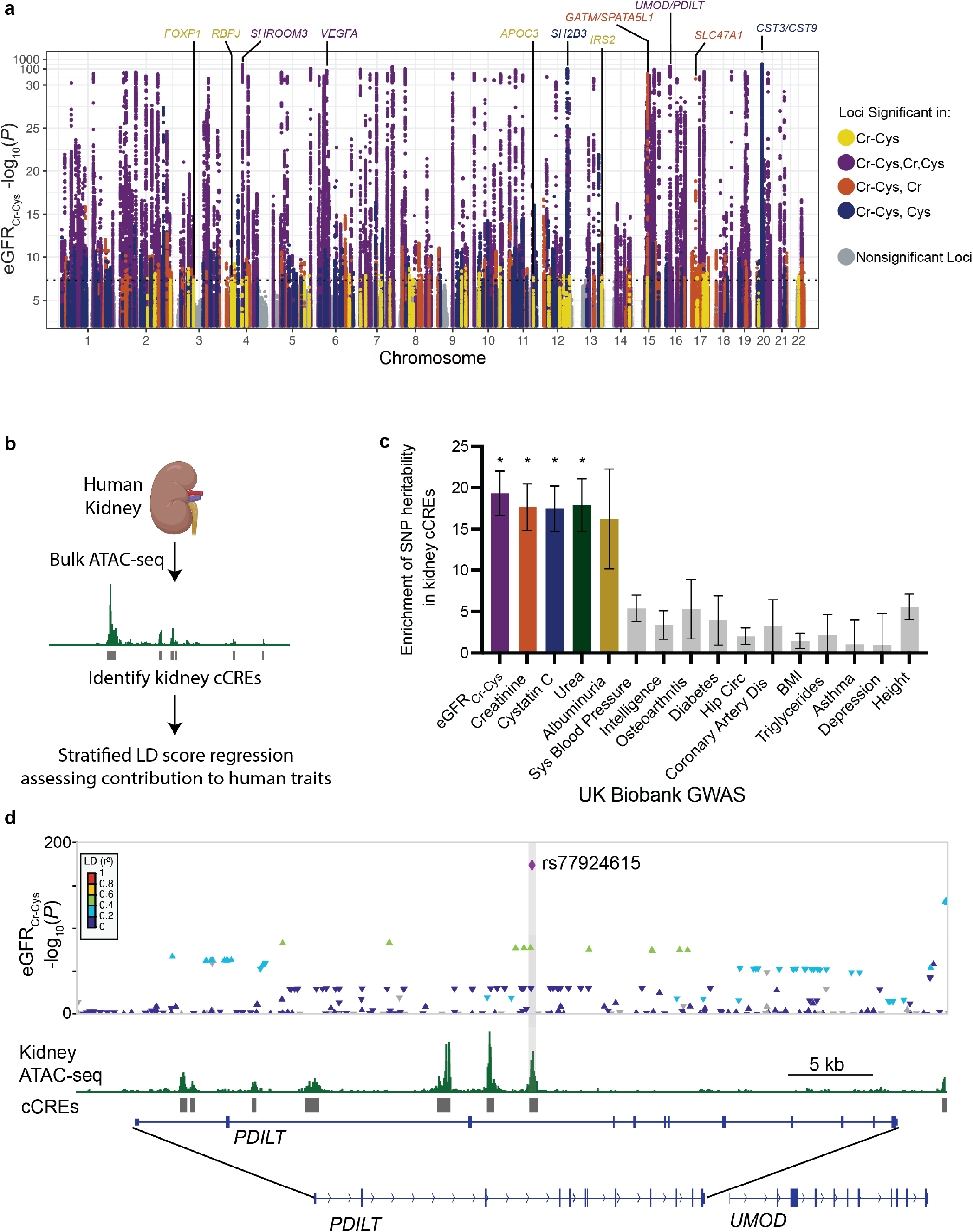
Kidney candidate *cis*-regulatory elements are enriched for kidney function heritability. **a)** Manhattan plot for the estimated GFR calculated from serum creatinine and cystatin C levels (eGFR_Cr-Cys_) genome wide association study (GWAS) performed within the UK Biobank. Significant loci are colored to indicate whether they are unique to the eGFR_Cr-Cys_ GWAS or are found within indicated single biomarker GWAS. Labels at the top indicate the closest gene to the index variant. The y-axis is a -log_10_(*P*) scale up to 30, after which it switches to a -log_10_[-log_10_(*P*)] scale. **b)** ATAC-seq of human kidney cortex identifies kidney noncoding candidate *cis*-regulatory elements (cCREs). The contribution of kidney cCREs to human traits is evaluated by calculating trait heritability cCRE enrichment using stratified LD score regression. **c)** Fold enrichment of the SNP heritability of fifteen human traits within kidney cCREs calculated with stratified LD score regression. Error bars denote jackknife standard errors for the enrichment estimate. These errors were used to calculate *P* values, which were Bonferroni corrected. * *P*<0.05. **d)** Several kidney cCREs lie in introns of *PDILT* 5’ to *UMOD*, which encodes a mendelian kidney disease gene. rs77924615 is associated with kidney function (eGFR_Cr-Cys_), is in a kidney cCRE, and has low LD with all other variants at the locus. Degree of LD to rs77924615 is indicated by variant color.

We compared the effect of index variants on eGFR_cr-cys_ to their effect on chronic kidney disease, as assessed by a GWAS performed in a distinct cohort^6^. Despite the chronic kidney disease GWAS having less power than the kidney function GWAS, the effects of variants on eGFR_cr-cys_ and chronic kidney disease were highly correlated (r=-0.69, p < 10^−64^) and 88% of the eGFR_cr-cys_ index variant alleles associated with decreased kidney function were associated with increased chronic kidney disease risk (Supplementary Figure 1a). Thus, kidney function variants affect chronic kidney disease risk.

We hypothesized that many genetic variants that affect kidney function alter kidney regulatory elements. To identify human kidney candidate *cis*-regulatory elements (cCREs), we performed an assay for transposase-accessible chromatin with sequencing (ATAC-seq) of human kidney cortex (Figure 1b). We identified over 190,000 regions of open chromatin, with a mean size of 523 bp, which we define as kidney cCREs. Kidney cCREs were strongly enriched for motifs of kidney transcription factors, such as HNF4A, HNF1B and TFCP2l1 (Supplementary Figure 1b). Using stratified linkage disequilibrium (LD) score regression of GWAS summary statistics^22^, we evaluated whether kidney cCREs contained variants which explain heritable differences in kidney function. Kidney cCREs were strongly enriched for kidney function variants affecting eGFR_Cr-Cys_ (as well as variants affecting individual biomarkers of kidney function including creatinine, cystatin C, and blood urea nitrogen levels, Figure 1c).

To better understand the relationship between genetic variants and kidney cCREs, we focused on loci where LD structure makes it possible to nominate a single variant as causal. One such locus included *UMOD*. Mutations in *UMOD* cause autosomal dominant tubulointerstitial kidney disease and variants proximal to *UMOD* have been associated with CKD and kidney function in multiple studies^6,7,9^. rs77924615 is a variant 5’ of *UMOD* that is strongly associated with kidney function and has low LD (r^2^ < 0.6) with any other variant in European populations. Notably, this variant falls within a kidney cCRE (Figure 1d). As additional examples, other individual putative causal variants identified from LD structure fell in kidney cCREs close to *DDX1* as well as *TFEB* and *HOXD8,* genes with known roles in kidney development and function Supplemental Figure 2a-c)^23,24^. Together, these findings suggest that many genetic variants that influence kidney function act by perturbing kidney cCREs.

### Identification of kidney cell type-specific regulatory elements

Genetic variants affecting kidney function could potentially affect the function of many cell types. As we found that variants affecting kidney function are highly enriched in kidney cCREs and many cCREs are cell type-specific, we reasoned that identification of cell type-specific cCREs could help identify which cell types are affected by kidney function variants. Therefore, we identified cCREs for each human kidney cell type using single-cell ATAC-seq (scATAC-seq).

We performed scATAC-seq of cortex and medulla from three donors to construct a chromatin accessibility atlas of the human kidney. After quality control, we profiled 34,240 cells, which associated into 10 clusters. To identify the cell types present in the scATAC-seq data, we used gene activity scores, a measure of aggregate chromatin accessibility around each gene that correlates with gene expression^25^. By correlating gene activity scores and gene expression, we integrated the scATAC-seq data with single-cell RNA sequencing (scRNA-seq) data^26^. This correlation allowed us to assign the scATAC-seq clusters to the following cell types: podocytes, parietal epithelial cells, proximal tubule, loop of Henle, distal tubule, collecting duct intercalating cells, endothelial cells, stromal cells, T cells and other immune cells (Figure 2a,b).

**Figure 2:**
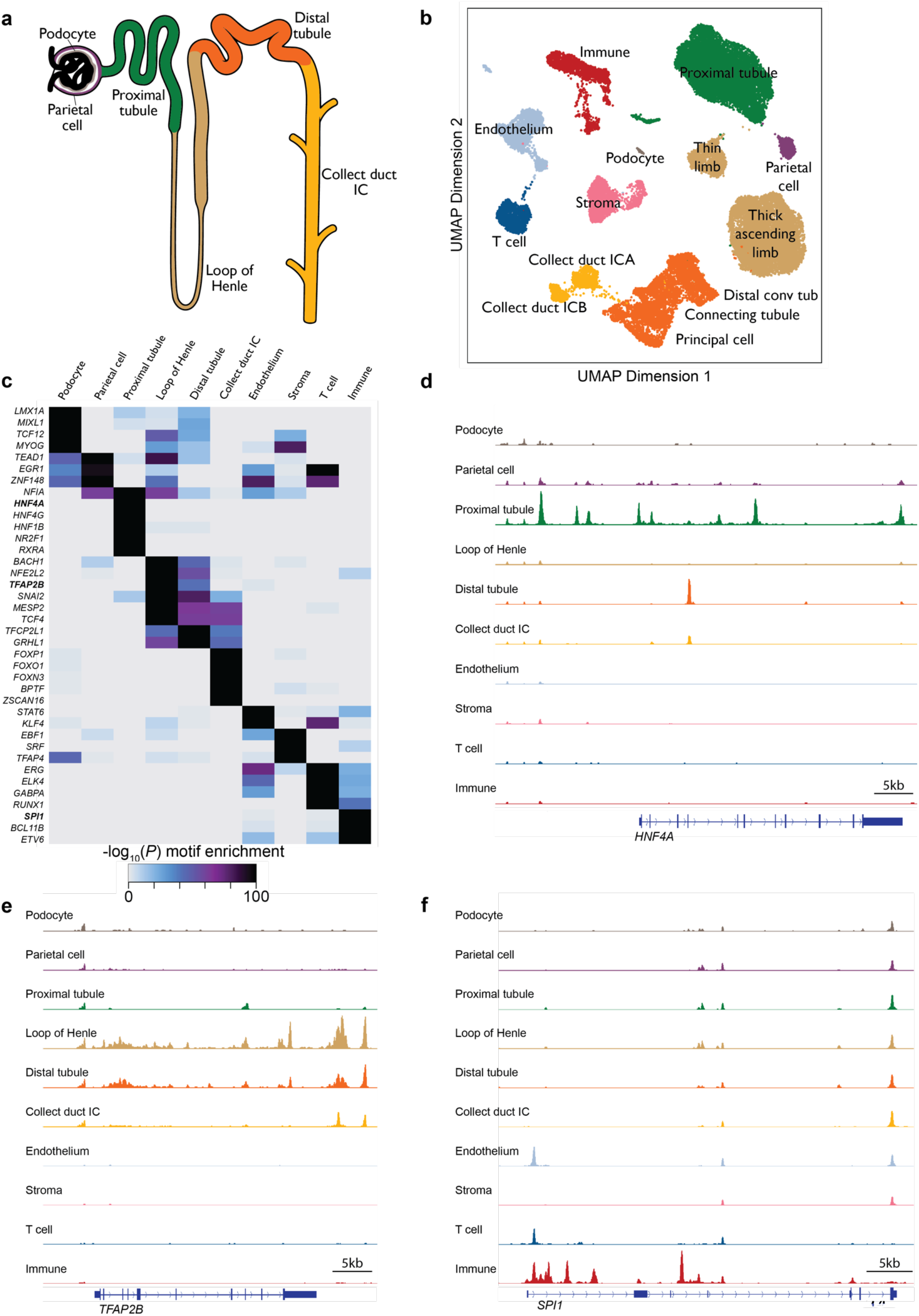
Single-cell ATAC-seq of human kidneys identifies cell type-specific cCREs. **a)** Schematic of the nephron, colored by major epithelial cell types. **b)** scATAC-seq UMAP of 34,240 kidney cells from 3 donors. **c)** Motifs of transcription factors expressed in the kidney are enriched in cCREs of specific kidney cell types. Chromatin accessibility for bolded transcription factors is shown in **d-f**. **d-f)** Chromatin accessibility maps for three cell type-specific genes: *HNF4A*, expressed in the proximal tubule; *TFAP2B*, expressed in the loop of Henle and distal tubule; and *SPI1*, expressed in immune cells.

To test the accuracy of cell type labels, we used gene activity scores to assess marker gene expression. High gene activity scores were observed for genes specifically expressed by cells of the corresponding clusters, including *NPHS1* (encoding Nephrin) in the podocyte cluster, *CUBN* (encoding Cubilin) in the proximal tubule cluster, *UMOD* (encoding Uromodulin) in the loop of Henle cluster*, FLT1* (encoding Vascular endothelial growth factor receptor 1) in the endothelial cell cluster, and *CD247* (encoding T-cell surface glycoprotein CD3ζ chain) in the T cell cluster (Supplementary Figure 3). Thus, gene activity scores of marker genes confirm that clusters of the kidney cCRE atlas are accurately assigned to cell types.

We assessed the enrichment of transcription factor binding motifs within cCREs of each cell type. The binding motif for HNF4A, a major transcriptional regulator of proximal tubule differentiation^27^, was enriched within cCREs accessible in proximal tubule cells (Figure 2c). Similarly, the binding motif for TFAP2B, which is necessary for differentiation of the distal tubule^28^, was enriched within cCREs accessible in distal tubule and loop of Henle cells (Figure 2c). And the binding motif for SPI1, necessary for immune cell differentiation^29^, was enriched for cCREs accessible in immune cells (Figure 2c).

Interestingly, multiple cell type-specific cCREs resided near genes encoding the transcription factors whose binding motifs were enriched in those same cell types. For instance, multiple cCREs accessible specifically in proximal tubule cells were near *HNF4A* (Figure 2d), cCREs accessible specifically in distal tubule and loop of Henle cells were near *TFAP2B* (Figure 2e), and cCREs accessible specifically in immune cells were near *SPI1* (Figure 2f). Thus, we observed both cell type-specific correlation of cCREs likely to mediate expression of transcriptional regulators and the cCREs through which those transcriptional regulators may function.

Genome-wide, we identified ∼500,000 cCREs, 15% of which were accessible in all assayed cell types (Figure 3a). However, most kidney cCREs were accessible in a single cell type or specifically in a few related cell types. For example, 9% of kidney cCREs were accessible in all tubule epithelial cells, but not other kidney cell types, another 8% were accessible specifically in proximal tubule epithelial cells, and another 12% were accessible specifically in distal tubule epithelial cell types (Figure 3a).

**Figure 3:**
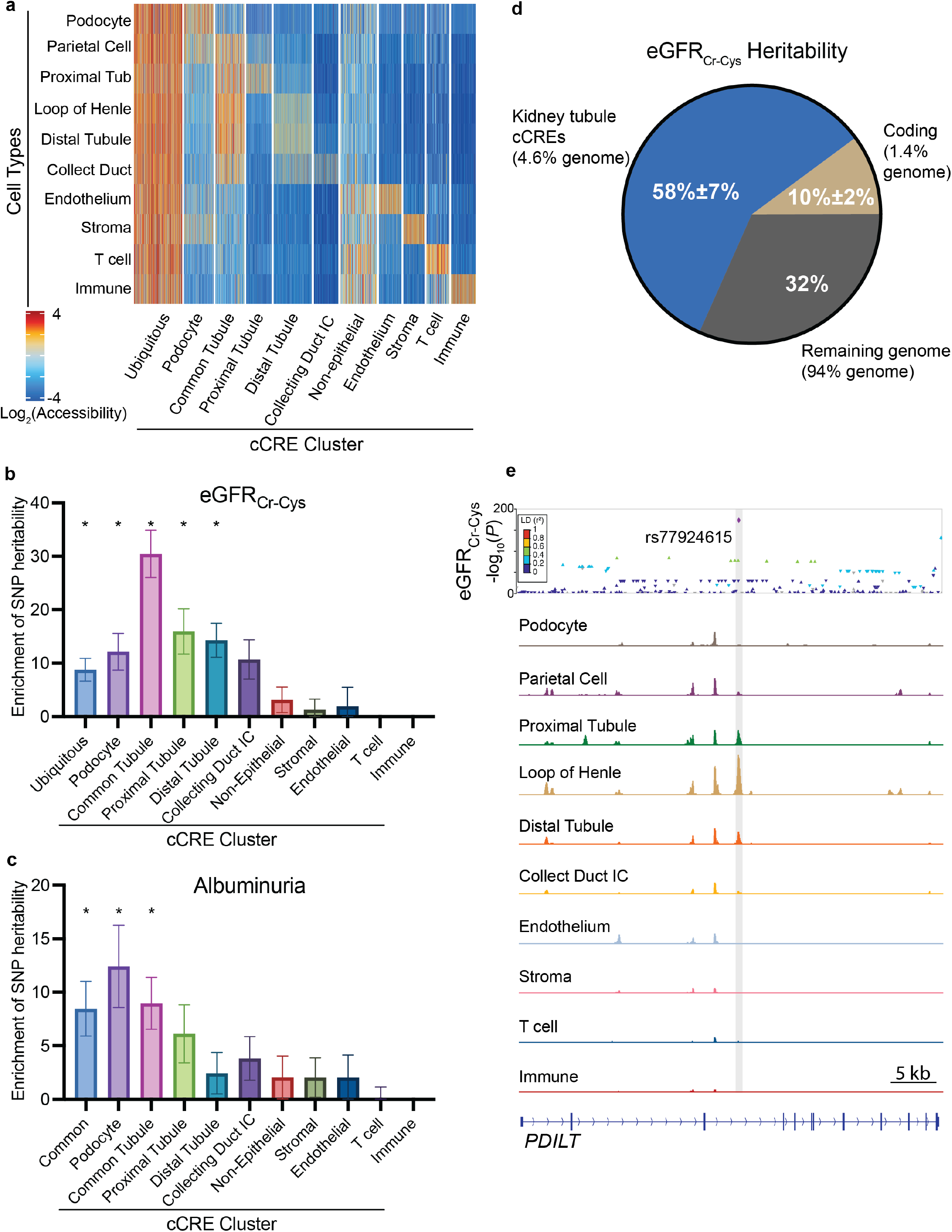
Kidney tubule cCREs account for the majority of SNP heritability of kidney function biomarkers. **a)** Clustering of cCREs distinguishes cell type-specific and ubiquitously accessible cCREs. Each vertical line represents one cCRE. A total of 526,273 cCREs are depicted. IC, intercalated cell. **b-c)** Fold enrichment of SNP heritability of eGFR_Cr-Cys_ and albuminuria within cell type-specific and ubiquitously accessible cCREs calculated with stratified LD score regression. Error bars denote jackknife standard errors for the enrichment estimate. These errors were used to calculate *P* values, which were Bonferroni corrected for multiple hypothesis testing. * *P*<0.05. **d)** Fraction of heritability of eGFR_Cr-Cys_ explained by kidney tubule epithelial cell cCREs (blue), coding exons (beige), and the remainder of the human genome (gray). **e)** rs77924615, a variant associated with eGFR_Cr-Cys_ level, lies with a tubule epithelial-specific cCRE (gray).

### Heritable differences in kidney function reflect differences in tubule epithelial cells and podocytes

To identify which cell types contribute to kidney function, we used stratified LD score regression to assess whether kidney function variants were enriched in cCREs of specific cell types (Figure 3b). Surprisingly, cCREs specifically accessible in endothelial cells, fibroblasts, or immune cells were not enriched for kidney function heritability. In marked contrast, cCREs accessible in specific kidney tubule epithelial cell types were 10-15-fold enriched for kidney function heritability and cCREs accessible in all kidney tubule epithelial cell types were 30-fold enriched for kidney function heritability. In addition, cCREs specific to podocytes were 12-fold enriched for kidney function heritability. These data implicate tubule epithelial cells and podocytes (and not endothelial cells, fibroblasts, or immune cells) as the major cell types affected by genetic variants that determine kidney function.

We also examined whether heritability of single kidney biomarkers (i.e., serum creatinine, blood urea nitrogen, and cystatin C levels) were enriched in cCREs of specific cell types (Supplementary Figure 4a-c). Serum creatinine level is affected by proximal tubule secretion of creatinine and its heritability was more strongly enriched in proximal tubule-specific cCREs than was blood urea nitrogen or cystatin C heritability. In contrast to creatinine, blood urea nitrogen level is regulated by both kidney function and reabsorption in the distal tubule. Consistent with this additional dependence on distal tubule function, blood urea nitrogen heritability was more strongly enriched in distal tubule-specific cCREs than was serum creatinine level heritability.

Podocytes form the filtration barrier that excludes large proteins from the urine^30^. In contrast to kidney function heritability, heritability of the marker of glomerular dysfunction—urine albumin—was most enriched in podocyte-specific cCREs (Figure 3c). We also noted enrichment of urine albumin heritability in tubule-specific cCREs, which we hypothesize is related to tubule reabsorption of filtered albumin^31^. Thus, the differences in the enrichment of blood urea nitrogen, serum creatinine and urine albumin-associated heritability were consistent with the specific biology of these biomarkers. These analyses further suggest that variants within regulatory elements affect kidney function through specific cell types and underscores the value of assaying kidney function using eGFR_Cr-Cys_ rather than single biomarkers.

To understand what fraction of kidney function heritability localized to kidney tubule epithelial regulatory elements, we used stratified LD score regression^22^. More specifically, we assessed the fraction of heritability of eGFR_cr-cys_ attributable to variants in cCREs accessible in the proximal tubule, loop of Henle, distal tubule or collecting duct. We found that variants in kidney tubule epithelial cell cCREs account for ∼58% of kidney function heritability (Figure 3d). Coding region variants accounted for an additional ∼10% of kidney function heritability. Thus, tubule epithelial cCREs and coding regions account for most kidney function common variant heritability. Podocyte specific-cCREs were also enriched for kidney function heritability; addition of podocyte cCREs to tubule epithelial cCREs and coding variants increased the explained fraction of kidney function heritability to 75% (Supplementary Figure 4d). Together, these data demonstrate that the majority of kidney function SNP-heritability lies in tubule epithelial cCREs, with smaller contributions from coding regions and podocyte cCREs.

To complement the stratified LD score regression analysis of kidney function heritability, we analyzed the likely causal variants identified from LD structure, discussed above. Most of these likely causal variants were found in tubule-specific cCREs. For instance, the *UMOD*-associated variant rs77924615 fell within a tubule-specific cCRE (Figure 3e).

### Distinguishing variants affecting kidney function from those affecting biomarker metabolism

We hypothesized that some loci identified in the eGFR_cr-cys_ GWAS reflected, not kidney function, but effects on individual biomarkers. For example, *CST3*, detected in the eGFR_cr-cys_ GWAS (Figure 1a), encodes cystatin C, one input into eGFR_cr-cys_. Therefore, it is likely that the effect of the *CST3* locus on eGFR_cr-cys_ is mediated by variants affecting cystatin C expression rather than kidney function (Figure 1A, Supplementary Table 1). In support of this possibility, the *CST3* locus is not associated with eGFR estimated from creatinine alone (Figure 1A, Supplementary Table 1).

We reasoned that loci which affect both creatinine and cystatin C are more likely to affect kidney function than loci which affect only a single biomarker^6^. Therefore, we assessed the effect of each eGFR_cr-cys_ index variant on eGFR estimated by either serum creatinine or cystatin C levels individually (Figure 4a). We identified 385 index variants which had the same direction of effect on serum creatinine and cystatin C and had significant effects (Bonferroni corrected p-value<0.05) on both biomarkers (Figure 4b, Supplementary Table 2). We refer to these variants as kidney function index variants. For example, index variants near *UMOD* and *SOX9* have consistent effects on both creatinine and cystatin C and these loci therefore likely affect kidney function. This possibility is bolstered by prior work demonstrating that *UMOD* is a monogenic kidney disease gene and *SOX9* is a critical regulator of kidney development and injury response^32–34^.

**Figure 4:**
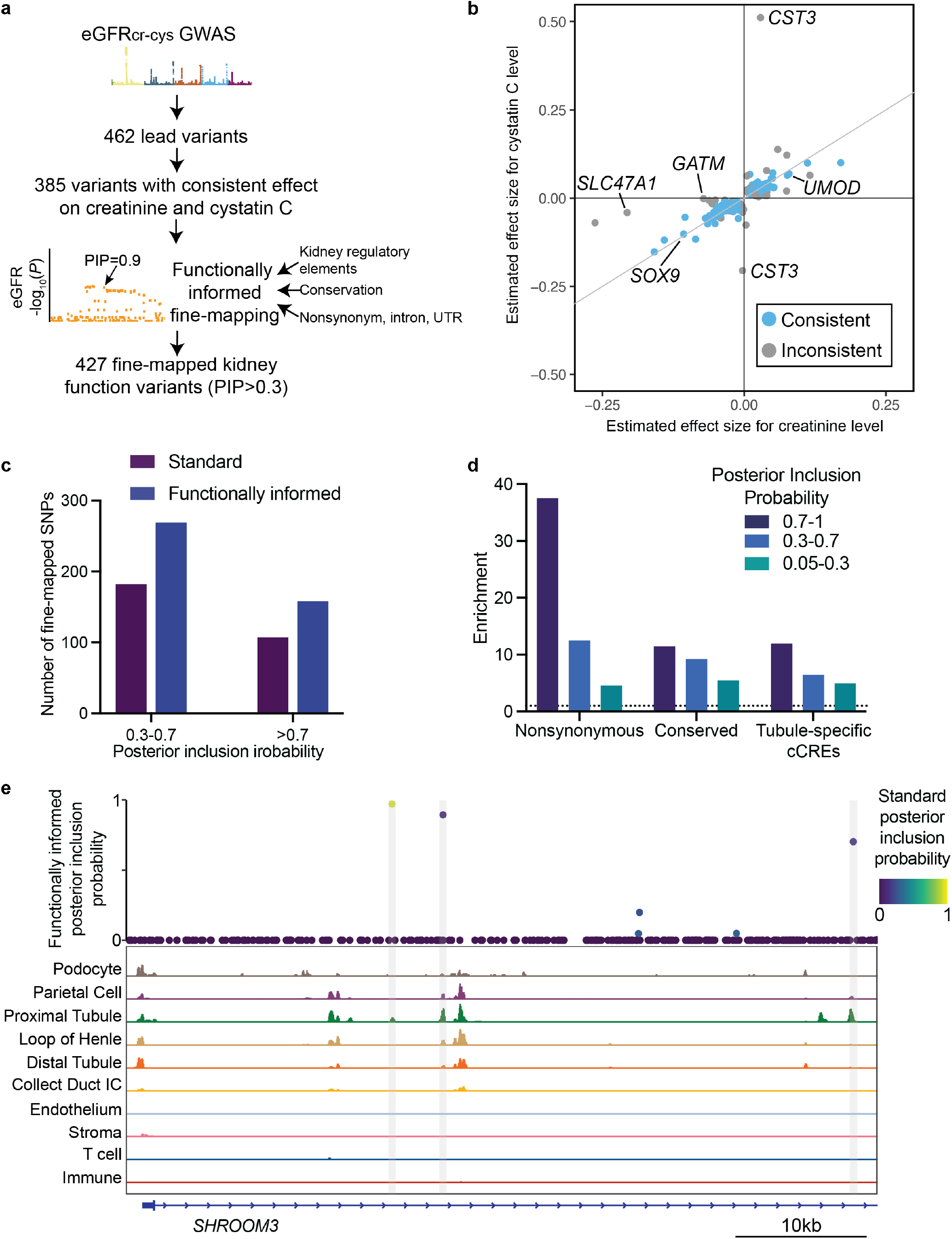
Identification of causal kidney function variants using functionally informed fine-mapping. **a)** Workflow of functionally-informed fine-mapping of variants affecting eGFR_cr-cys_. GWAS for eGFRcr-cys identified 462 lead variants. 385 of these variants had consistent effects on both serum creatinine and cystatin C levels. Fine-mapping of these loci incorporated annotation of kidney tubule cCREs, evolutionary conservation, and gene annotations including annotations indicating whether a variant is coding, affects protein sequence (nonsynonymous), is in an intron, or is in a UTR. PIP, posterior inclusion probability. **b)** eGFR_cr-cys_ index variants plotted by the estimated size of their effect on both serum creatinine and cystatin C levels. Variants colored blue affect both creatinine and cystatin C levels in the same direction. **c)** Comparison of the number of fine-mapped kidney function variants identified with or without functional annotations. Variants are separated into those with moderate posterior inclusion probability (0.3-0.7) and high posterior inclusion probability (>0.7). **d)** Fold enrichment of fine-mapped kidney function variants that cause nonsynonymous changes in a coding sequence, are evolutionarily conserved, or that lie within tubule epithelial-specific cCREs. Variants are separated into those with high (>0.7), moderate (0.3-0.7) and low (0.05-0.3) posterior inclusion probabilities. **e)** Functionally informed fine-mapping at the *SHROOM3* locus. The functionally informed posterior causal probability is plotted for variants, which are colored according to the causal probability estimated without functional annotations. Chromatin accessibility from scATAC-seq for the indicated kidney cell types are plotted below. Gray boxes indicate tubule epithelial cCREs containing fine-mapped variants.

77 index variants including 2 index variants near *CST3* affected only one biomarker (i.e., Bonferroni corrected p-value>0.05 for one biomarker) or exhibited discordant directions of effect (Figure 4b, Supplementary Table 3). Among index variants associated only with serum creatinine levels, several were at loci with plausible connections to known non-GFR determinants of serum creatinine levels. For instance, an index variant associated only with serum creatinine levels occurred in *GATM,* which encodes the rate-limiting enzyme in creatinine synthesis^35^. Similarly, an index variant associated only with serum creatinine levels occurred in *SLC47A1,* which participates in creatinine secretion^36,37^. Thus, determining which loci affect both creatinine and cystatin C is likely to distinguish loci that affect kidney function from those that affect biomarker metabolism.

### Functionally informed fine-mapping of kidney function variants

Within the GWAS-identified loci, which genetic variants affect kidney function? To begin to identify the causal variants, we applied functionally informed fine-mapping using PolyFun^38^ to 1 Mb centered around each kidney function index variant. PolyFun is a fine-mapping approach that can incorporate multiple genome-wide functional annotations to inform prior causal probabilities and improve fine-mapping power. We performed fine-mapping both with and without annotation of conservation, genes, as well as the kidney tubule epithelial cell and podocyte cCREs that we had defined. Inclusion of these functional annotations in fine-mapping identified 65 more variants with posterior inclusion probability >0.7 and 138 more variants with a posterior inclusion probability >0.3, corresponding to a 47% improvement in the number of fine-mapped variants relative to standard fine-mapping (Figure 4c). These fine-mapped variants were highly enriched for missense variants, conservation, and presence in tubule epithelial-specific cCREs (Figure 4d and Supplementary Figure 5a). For instance, at the *SHROOM3* locus^9^, three fine-mapped variants (PIP>0.4) occurred within tubule epithelial-specific cCREs (Figure 4e).

### Identification of candidate genes involved in human kidney function

Many of these fine-mapped variants implicate specific genes in human kidney function. 55 of these variants (PIP>0.4) were missense variants within coding sequence. These fine-mapped missense variants implicate 50 genes in kidney function (Supplementary Table 4). Ten of these

50 genes are associated with monogenic kidney phenotypes in humans^39–49^ (Table 1). An additional seven of these genes cause kidney phenotypes when mutated in model organisms^50–56^ (Table 1). Many of the genes were associated with primary cilia, regulation of mTOR, glucose metabolism, and regulation of calcium and phosphorous, biological processes with recognized links to kidney pathology^57–59^.

**Table 1:** Fine-mapped kidney function variants implicate genes involved in kidney function. A subset of genes with missense or promoter fine-mapped kidney function variants (PIP>0.4). Additional genetic evidence from the role of these genes in human monogenic kidney diseases or model organisms is noted. AD, autosomal dominant; AR, autosomal recessive.

| Gene | Additional Genetic Evidence | Function |
| --- | --- | --- |
| *ATP6V1B1* | AR Distal renal tubular acidosis and sensorineural hearing loss | Calcium/Phosphorous |
| *SLC34A3* | AR Hereditary hypophosphatemic rickets with hypercalciuria | Calcium/Phosphorous |
| *CLUAP1* | Zebrafish: Kidney cyst | Cilia |
| *PKHD1* | AR Polycystic kidney disease | Cilia |
| *POC5* |  | Cilia |
| *PKD1* | AD Polycystic kidney disease | Cilia |
| *EPB41L5* | Mouse: proteinuria, kidney failure | Extracellular matrix |
| *COL18A1* |  | Extracellular matrix |
| *ELN* |  | Extracellular matrix |
| *ALDOB* | AR Fructose intolerance | Glucose metabolism |
| *GCKR* |  | Glucose metabolism |
| *APOE* | AD Lipoprotein glomerulopathy | Lipid metabolism |
| *DEPTOR* | Mouse: Reduced kidney injury | mTOR |
| *FNIP1* | Mouse: Kidney cysts | mTOR |
| *LRP2* | AR Donnai-Barrow syndrome | Proximal tubule reabsorption |
| *SLC7A9* | AD/AR Cystinuria | Proximal tubule reabsorption |
| *KCNK5* | Mouse: Kidney fibrosis | Proximal tubule reabsorption |
| *GATA5* | Mouse: Kidney immune infiltrate | Transcription factor |
| *HNF1B* | AD Tubulointerstitial kidney disease | Transcription factor |
| *HNF4A* | AD Fanconi renal tubular syndrome | Transcription factor |
| *NFAT5* | Mouse: Kidney medulla atrophy | Transcription factor |

To identify target genes whose expression may depend on noncoding fine-mapped variants, we evaluated a nearest gene approach, a gene similarity-based approach—the Polygenic Priority Score (PoPS)^60^, and an integration of these two orthogonal approaches.

To assess the performance of these gene prioritization strategies for kidney function, we constructed gold-standard kidney function gene sets comprised of: 1) genes containing rare variants detected by exome sequencing in the UK Biobank (Supplementary Table 5) and 2) genes containing a high-confidence (PIP>0.7) fine-mapped missense variant (Supplementary Table 4). Using these gold-standard gene sets, we evaluated three gene prioritization strategies: 1) the gene with the top PoPS score within 500kb of the variant, 2) the nearest gene, and 3) integration of PoPS and the nearest gene, in which a gene is prioritized only if the top PoPS gene and nearest gene are the same. The nearest gene approach had a precision and recall of over 60%, PoPS had a precision and recall of 40-60%, and combining the PoPS and nearest gene approach gave a precision of 100% with a recall from 30-50% (Supplementary Figure 5b,c). As combining the PoPS score with analysis of the nearest gene identified targets more precisely than competing GWAS-based gene prioritization methods^60^, we used this integrated approach to identify a set of 67 genes that may be regulated by 80 noncoding fine-mapped kidney function variants (Supplementary Table 6). In total, we identified 111 putative kidney function genes by integrating the 67 predicted targets of fine-mapped noncoding variants with the 50 genes containing fine-mapped missense variants (Supplementary Table 7).

Gene enrichment analysis of kidney function genes revealed enrichment for tube development (*P*_adj_<10^−9^) and epithelium development (*P*_adj_<10^−8^) (Supplementary Table 8). In addition, genes encoding transcriptional regulators were enriched (*P*_adj_ <10^−11^), including a subset with previously described roles in kidney development (e.g., *HNF4A, MECOM, TFAP2B, TFCP2L1,* and *NFIA*) and tubule epithelial response to stress (e.g., *FOXO3* and *NFATC1*) ^27,61–68^. Specifically enriched pathways included AMPK signaling (*P*_adj_ <10^−2^) and HIF-1 signaling (*P*_adj_ <10^−2^). Together, these results highlight the importance of genes involved in tubule epithelial cell development and response to tubule stress for variation in human kidney function.

### Kidney function variants affect tubule epithelial chromatin accessibility

As the genetic variants within kidney tubule epithelial cCREs explain ∼58% of eGFR_cr-cys_ SNP-heritability (Figure 3d), we sought to assess how variants in these cCREs might affect kidney function. One potential mechanism is altering tubule gene expression by changing chromatin at regulatory elements. ATAC-seq can detect the effect of variants on local chromatin accessibility^69^. To identify variants that affect chromatin accessibility, we performed allele-specific mapping of our scATAC-seq data and identified heterozygous variants at which there was a significant difference in accessibility between the reference and alternate allele.^70^ We refer here to such variants as displaying chromatin accessibility allelic imbalance (CAAI). 2,305 variants displayed CAAI in proximal tubule cells, 1,251 variants displayed CAAI in loop of Henle cells, and 5,852 variants displayed CAAI in jointly analyzed tubule epithelial cells (FDR <0.05, Figure 5a).

**Figure 5:**
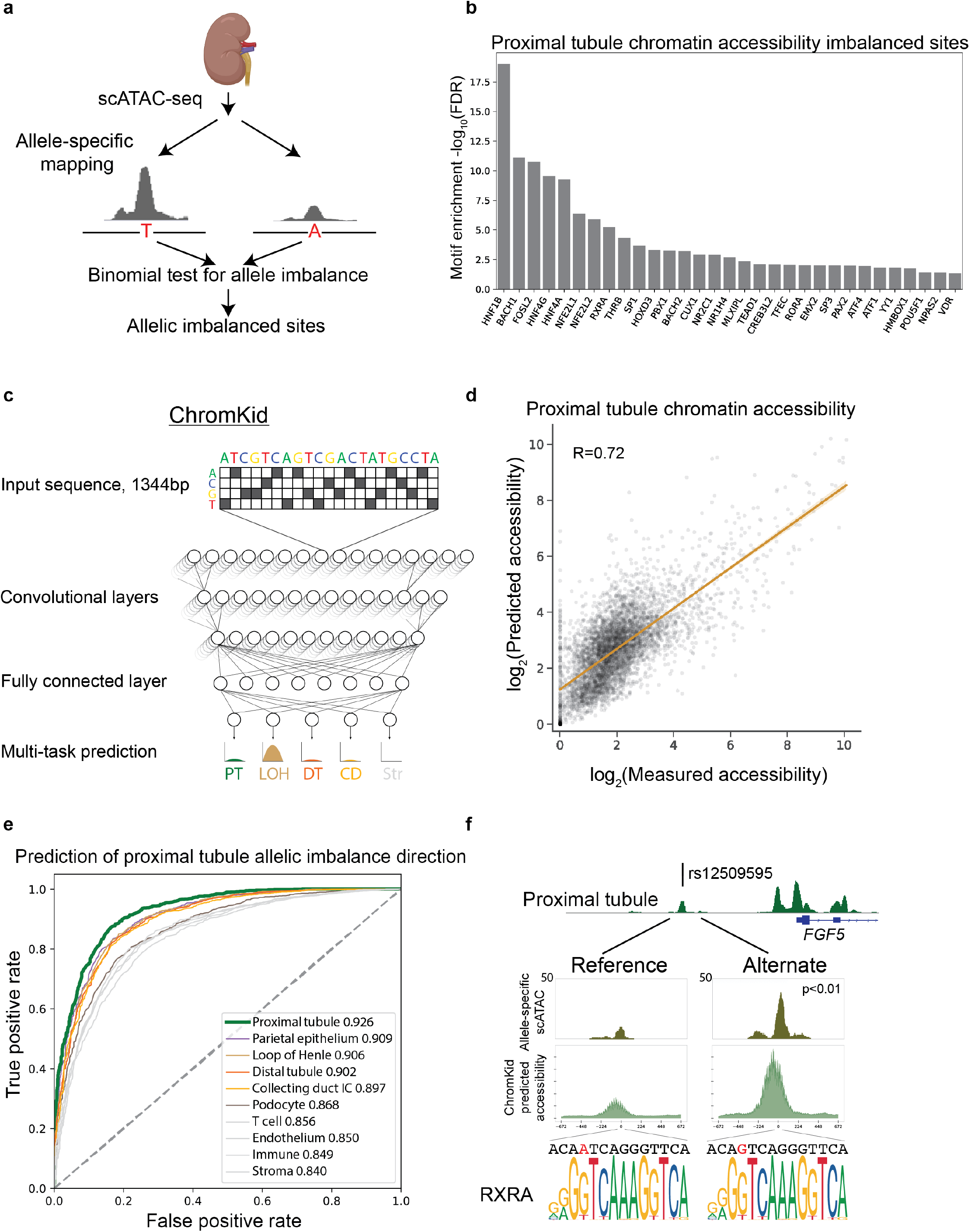
Chromatin accessibility allelic imbalance and machine learning identify cell type-specific effects of variants on chromatin accessibility. **a)** Workflow to detect chromatin accessibility allelic imbalance (CAAI). Single-cell ATAC-seq reads were analyzed at heterozygous alleles and accessibility at both alleles was compared to measure the effect of variants on chromatin accessibility. **b)** Enrichment of proximal tubule expressed transcription factor-binding motifs at cCREs exhibiting CAAI in the proximal tubule. **c)** Schematic of ChromKid, a convolutional neural network trained to predict cell type-specific chromatin accessibility for 10 kidney cell types from DNA sequence. ChromKid uses a 1344bp DNA sequence input to generate a prediction of the quantitative chromatin accessibility at the center of the input region for each kidney cell type. **d)** Measured versus ChromKid-predicted proximal tubule chromatin accessibility on chromosome 11, which was withheld from training. The best fit line is shown. **e)** Receiver operating characteristic (ROC) curves for the direction of proximal tubule chromatin accessibility imbalance, displaying the false positive rate (x-axis) versus true positive rate (y-axis) of cell-type specific predictions from ChromKid. The areas under the curves (AUCs) for ChromKid predictions for each cell type are shown in the legend. ChromKid predictions of variant effects in each cell type are compared to proximal tubule CAAI. **f)** Fine-mapped variant rs12509595 (indicated by a vertical line) maps to a proximal tubule cCRE near *FGF5*. CAAI analysis revealed the cCRE with the alternate allele exhibits increased chromatin accessibility in the proximal tubule. ChromKid also predicted that the alternate allele would show increased chromatin accessibility at this cCRE in the proximal tubule. A motif in this cCRE corresponds to a consensus binding motif for the proximal tubule transcription factor RXRA. The variant, indicated in red, affects this binding motif. The alternative allele sequence more closely matches the consensus binding motif for RXRA. Statistical significance was calculated with a two-sided binomial test.

We hypothesized that a mechanism by which variants affect chromatin accessibility is by perturbing the binding of transcription factors. Consistent with this hypothesis, CAAI sites were enriched in transcription factor binding motifs of transcription factors expressed in the relevant cell type (Figure 5b). For instance, proximal tubule CAAI sites were enriched for binding motifs for HNF1B and HNF4A, two proximal tubule transcriptional regulators.

An additional hypothesis is that transcriptional activators increase the chromatin accessibility of alleles which more closely match their consensus binding motifs (Supplementary Fig 6a). As hypothesized, alleles that more closely approximated HNF1B and HNF4A binding motifs were associated with increased chromatin accessibility in proximal tubule cells (Supplementary Fig 6b). In contrast, alleles that more closely match the consensus transcription factor binding motif of transcriptional repressors CUX1 and YY1 were associated with less chromatin accessibility (Supplementary Fig 6b). Thus, CAAI analysis of scATAC-seq data can reveal how variants affect regulatory elements within specific cell types and implicate specific transcriptional regulators in their function.

A limitation of CAAI analysis is that it depends on the presence of heterozygous alleles represented in the scATAC-seq data. In an attempt to overcome this hurdle, we developed ChromKid, a convolutional neural network (CNN), to predict chromatin accessibility in each kidney cell type from DNA sequence (Figure 5c). More specifically, ChromKid is a multitask CNN trained on our kidney scATAC-seq data to predict chromatin accessibility in each of 10 kidney cell types. ChromKid’s predictions of chromatin accessibility correlated with measured accessibility (R=0.73 for proximal tubule, Figure 5d). The strength of the correlation between predicted and measured chromatin accessibility varied depending on kidney cell type, with the highest correlation in tubule epithelial cell types and lowest correlation in podocytes (Supplementary Figure 6c). These differences in degree of correlation were explained by differences in the number of cells present in the training data for each cell type (Supplementary Figure 6c).

Since ChromKid only uses DNA sequence to predict chromatin accessibility, it can predict the effect of any variant on local chromatin accessibility. To assess the accuracy of ChromKid variant effect predictions, we compared ChromKid predictions to experimental CAAI data. ChromKid predictions of variant effects in proximal tubule accurately discriminated whether a variant increased or decreased proximal tubule chromatin accessibility (AUROC 0.93, Figure 5e). ChromKid’s proximal tubule predictions more accurately discriminated variants effects on proximal tubule chromatin accessibility than ChromKid’s predictions for other kidney cell types (AUC 0.84-0.91), with the degree of inaccuracy correlating with the distance of the lineage relationship to proximal tubule cells (Figure 5e). Thus, ChromKid learns cell type-specific information regarding how variants affect chromatin accessibility.

In addition to predicting whether variants increased or decreased chromatin accessibility, we assessed whether ChromKid could distinguish which variants affect chromatin accessibility. We used the experimental CAAI data to identify sets of variants that did or did not affect chromatin accessibility in proximal tubule cells and assessed whether ChromKid could accurately distinguish CAAI variants. Indeed, ChromKid identified many of the variants that affect proximal tubule chromatin accessibility (AUROC 0.77, Supplementary Figure 6d). ChromKid’s proximal tubule predictions more accurately discriminated which variants affect proximal tubule chromatin accessibility than ChromKid’s predictions for other kidney cell types, again with the degree of inaccuracy correlating with the distance of the lineage relationship (Supplementary Figure 6d). Together, these data indicate that ChromKid predicts how genetic variants affect chromatin accessibility in different cell types.

To begin to test the possibility that CAAI and ChromKid may help identify how noncoding genetic variants affect kidney function, we examined rs12509595, a fine-mapped variant for eGFR_cr-cys_. rs12509595 is in a proximal tubule cCRE near *FGF5*. This variant displayed CAAI in proximal tubule cells, with the alternate allele exhibiting more accessibility than the reference allele (Figure 5f). Correspondingly, ChromKid predicted that the alternate allele would increase chromatin accessibility in proximal tubule cells (Figure 5f). The concordance of the measured and predicted effects on chromatin accessibility increased confidence that this variant (and not a distinct linked variant) was responsible for the effect on chromatin accessibility.

Can CAAI and ChromKid also help illuminate how rs12509595 might affect *FGF5* expression? RXRA is a transcription factor expressed in the proximal tubule and helps regulate response to kidney injury^71,72^. We found that RXRA-binding motifs are enriched at CAAI sites, including that containing rs12509595 (Figure 5b). Within RXRA-binding motifs, the allele that better matched the RXRA-binding consensus sequence was associated with increased chromatin accessibility in proximal tubule cells (Supplementary Figure 6b). The alternate rs12509595 allele more closely matches the RXRA-binding consensus, suggesting that this allele may increase chromatin accessibility in proximal tubule cells by increasing binding to RXRA. This example illustrates how CAAI and ChromKid can identify mechanisms by which variants may affect chromatin architecture and kidney function.

### Kidney function genetic variants affect tubule epithelial regulatory element function

We hypothesized that many genetic variants affect kidney function by altering tubule epithelial regulatory element function. To test this hypothesis, we employed ChromKid to predict the effect of tubule epithelial cCRE variants on chromatin accessibility. Notably, fine-mapped tubule epithelial cCRE variants had larger predicted effects on chromatin accessibility than other tubule epithelial cCRE variants (Figure 6a), consistent with the proposal that many genetic variants affect kidney function by altering tubule epithelial chromatin accessibility.

**Figure 6:**
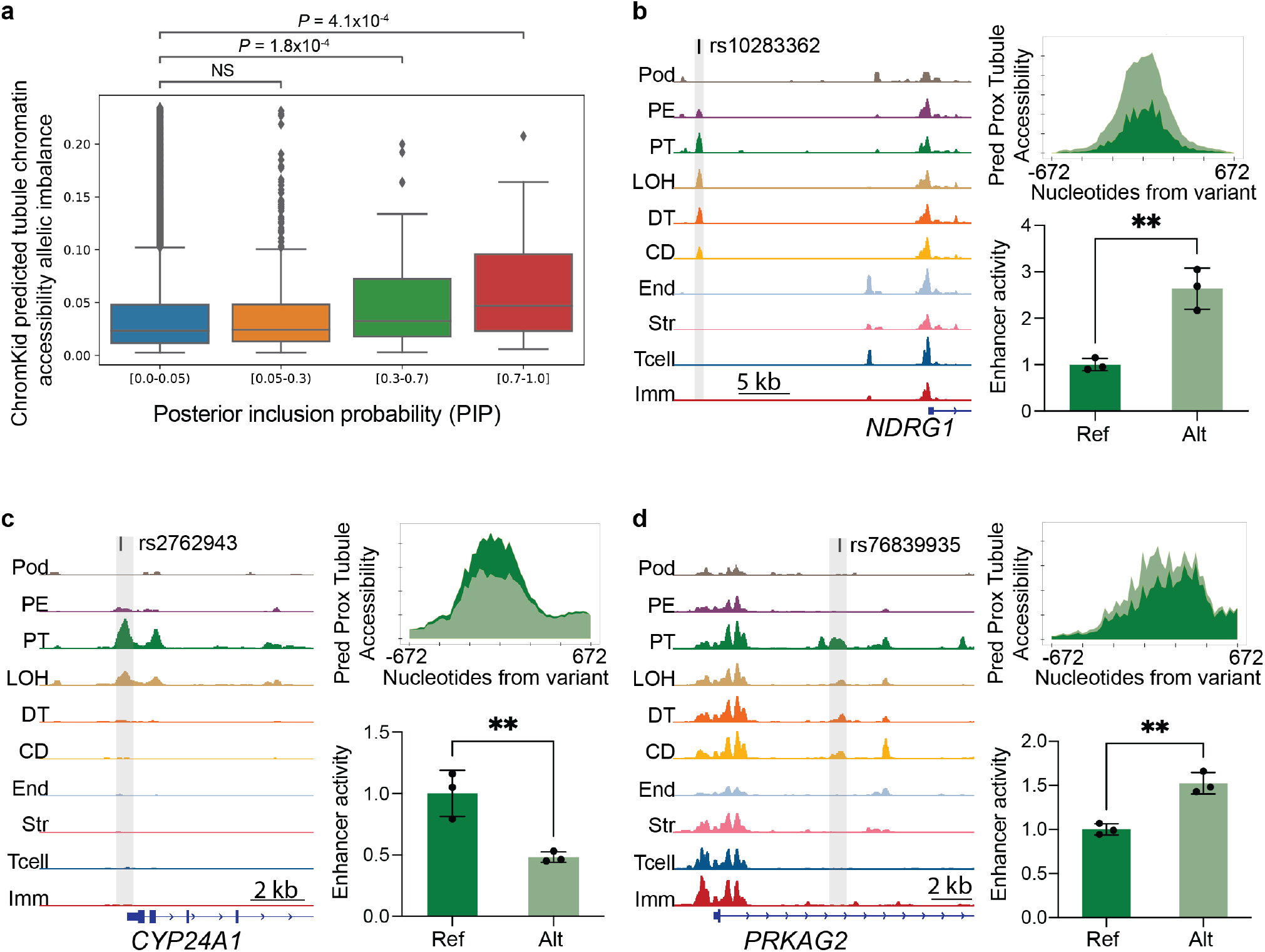
Genetic variants affecting kidney function alter tubule epithelial *cis*-regulatory element function. **a)** Fine-mapped kidney function variants exhibit larger predicted effects on accessibility than other variants within tubule epithelial cCREs. ChromKid-generated predictions of CAAI at fine-mapped kidney function (eGFR_cr-cys_) variants within tubule cCREs stratified by PIP. Predicted CAAI is shown as the 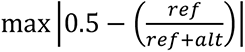 across tubule epithelial cell types. Box boundaries indicate the 1^st^ and 3^rd^ quartile and whiskers indicate the most extreme data point within 1.5 times the interquartile range. Statistical significance was calculated using the Mann Whitney U test. **b-d)** Left: Plots of scATAC-seq data near fine-mapped kidney function variants, which are depicted by vertical lines. cCREs (gray) containing a kidney function variant were selected for functional testing. Top right: ChromKid predicted proximal tubule chromatin accessibility for both alleles at these cCREs. Predicted chromatin accessibility for the reference allele is depicted in dark green, and for the alternative allele is depicted in light green. Bottom right: Activity of enhancers in human primary tubule epithelial cells. Enhancer activity of the reference allele is depicted in dark green, and of the alternative allele is depicted in light green. Results are representative of three independent experiments. Statistical significance was based on a two-sided *t* test, ** *P*<0.01.

To test whether kidney function variants alter tubule epithelial cell regulatory element function, we performed enhancer assays in tubule epithelial cells. More specifically, we measured the effects of fine-mapped kidney function variants on the activity of three cCREs (5’ of *NDRG1*, in the promoter of *CYP24A1*, and in an intron of *PRKAG2*) in cultured primary human tubule epithelial cells (PHTE). For the *NDRG1*-associated cCRE, the alternative allele of rs10283362 increased enhancer activity (Figure 6b). Similarly, the other fine-mapped variants affected the activity of their associated enhancers (Figure 6c,d). Moreover, the alleles with increased enhancer activity were also predicted by ChromKid to increase chromatin accessibility. Together, these data support the hypothesis that many variants affect kidney function via effects on tubule epithelial regulatory elements.

### CRISPRi enhancer-gene mapping links genes to kidney function

Linking variants and cCREs to the genes they regulate remains a major challenge. Measuring the effect of cCREs on gene expression can identify which genes those cCREs control. To begin to map cCREs harboring kidney function variants to kidney function genes, we deployed CRISPRi enhancer-gene mapping (Figure 7a).

**Figure 7:**
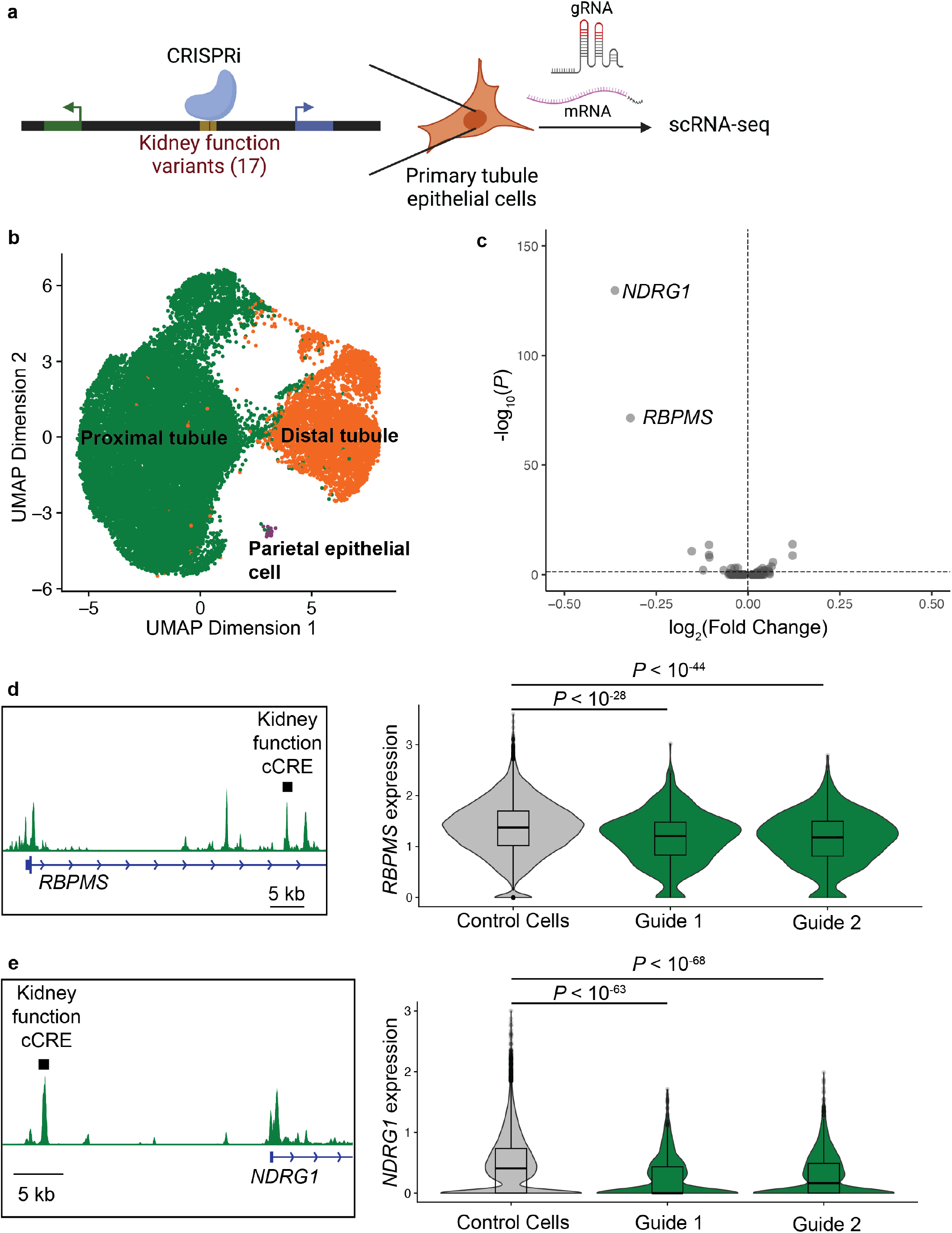
Kidney function enhancers regulate *RBPMS* and *NDRG1*. **a)** Schematic of the approach to map genes regulated by kidney function regulatory elements using CRISPRi-mediated silencing. cCREs containing fine-mapped kidney function variants were targeted in human primary tubule epithelial cells with CRISPRi. Gene expression and guide expression were measured by scRNA-seq. **b)** UMAP of scRNA-seq for 24,563 primary tubular epithelial cells from 4 donors used for CRISPRi-mediated silencing of kidney function cCREs. **c)** Volcano plot of differentially expressed genes in cells with expression of guides targeting kidney function cCREs. **d-e**). Left: Plots of proximal tubule chromatin accessibility from scATAC-seq data with the CRISPRi-targeted cCRE depicted with a black rectangle. Right: Violin plots of gene expression in cells expressing (green) or not expressing (gray) guides targeting the indicated regulatory element. Superimposed box plots indicate the median, 25^th^, and 75^th^ percentiles of expression. All *P* values are adjusted for multiple hypothesis testing using the Bonferroni correction.

Complicating the functional analysis of variants, most cCREs harboring fine-mapped kidney function variants were not accessible in immortalized kidney epithelial cell lines (Supplementary Figure 7a). Therefore, we assessed cCRE function in PHTEs (Figure 7a). We designed dual sgRNAs vectors targeting 17 tubule cCREs, each of which contained a fine-mapped kidney function variant. For positive and negative controls, we generated sgRNAs targeting 6 previously identified enhancers, 6 transcriptional start sites, 4 gene deserts, 4 regions of closed chromatin, and 4 nontargeting controls (Supplementary Table 9).

We identified cCRE target genes by measuring the effect of cCRE silencing on gene expression. More specifically, we used scRNA-seq to measure the effect of cCRE silencing on gene expression in over 24,000 HRCEs from four donors. scRNA-seq revealed that the HRCEs were primarily proximal tubule and distal tubule cells with a small number of parietal epithelial cells (Figure 7b and Supplementary Figure 7b). After lentiviral transduction with CRISPRi and sgRNAs, a median of 1,257 cells expressed each guide and these cells expressed a median of 3 guides (Supplementary Figure 7c).

All guides directed against previously identified enhancers^73^ and TSSs downregulated the associated gene, as expected (Supplementary Figure 7d,e). CRISPRi-mediated silencing of two kidney function-associated cCREs downregulated a nearby gene (Figure 7c). Guides directed against a cCRE containing a kidney function variant in the first intron of *RNA-binding protein with multiple splicing* (*RBPMS*) downregulated *RBPMS* (Figure 7c). Similarly, guides directed against a cCRE containing a kidney function variant 5’ of *N-Myc downstream regulated 1* (*NDRG1*) reduced *NDRG1* expression (Figure 7d). Effects were consistent across all 4 donors (Supplementary Figure 8a,b). In addition, the regulatory element which regulates *NDRG1*, contains rs10283362, the fine-mapped kidney function variant that affects tubule epithelial cell enhancer function (Figure 6b). These data indicate that variants in regulatory elements controlling *NDRG1* and *RBPMS* expression in tubule epithelial cells affect human kidney function.

## Discussion

To identify cellular and molecular mechanisms that affect kidney function in humans, we performed a GWAS of kidney function, identified likely causal variants, and determined how variants affect chromatin accessibility, regulatory element function and gene expression. The pipeline of GWAS, cCRE-informed fine mapping, enhancer assays and CRISPRi enhancer-gene mapping overcomes the challenges posed by linkage disequilibrium to reveal how non-coding variants affect the cells and genes involved in kidney function. Thus, in addition to identifying variants and genes involved in human kidney function, this work illustrates a strategy for moving from GWAS to variants, regulatory elements, and genes extensible to other polygenic traits.

Variants in kidney podocyte, proximal tubule, and distal tubule-specific cCREs were enriched for kidney function heritability. Strikingly, variants in tubule epithelial cell cCREs accounted for the majority of kidney function heritability. As kidney function and chronic kidney disease GWAS were highly correlated, we propose that differences in tubule epithelial cell biology are major drivers of human chronic kidney disease risk.

Heritability of serum creatinine level reveals a quantitatively larger role for the proximal tubule than other tubule cell types and prior studies have highlighted the involvement of the proximal tubule in kidney function heritability^12,13,74,75^. Because the proximal tubule actively secretes creatinine and therefore has a unique filtration-independent role in creatinine clearance, focus on serum creatinine as a sole measure of GFR likely overestimates the role of the proximal tubule in kidney function. Inclusion of kidney function biomarkers beyond creatinine including cystatin C and blood urea nitrogen revealed that heritability of kidney function is distributed between podocytes, proximal tubule-specific cCREs, distal tubule specific cCREs, and most strikingly in cCREs shared by all tubule epithelial cell types.

A corollary of the finding that differences in tubule epithelial cells drive differences in kidney function is that tubule-directed therapies may be useful for treating many types of kidney disease. Additional support for this possibility comes from inhibitors of sodium-glucose transporter 2 (SGLT2). Inhibitors of SGLT2, which reabsorbs glucose in the proximal tubule, are effective for a variety of kidney diseases^76,77^. Our genetic data are consistent with a central role for proximal tubule cells in diverse kidney diseases and suggest that therapies targeting other tubule epithelial cells or other aspects of proximal tubule biology may be clinically effective.

To dissect how variants affect regulatory elements, we measured how variants affect chromatin accessibility. We used allele-specific scATAC-seq data to measure how variants impact chromatin accessibility (CAAI). CAAI is limited to variants that are heterozygous in the profiled cells. Moreover, interpretation is complicated by the potential effect of other variants in the haplotype. To address these limitations, we developed ChromKid, a deep learning model for predicting kidney cell chromatin accessibility from DNA sequence. These complementary biological and computational approaches revealed that many kidney function variants affect chromatin accessibility in tubule epithelial cells.

Identification of the cells and regulatory elements affected by causal variants helps identify the genes affected by those variants^73,78,79^. For the fine-mapped variants identified in our study, we nominated associated genes using a combination of computational and experimental approaches. The computational methods integrated nearest-gene assessment and gene similarity-based predictions^60^. The experimental methods made use of enhancer assays and CRISPRi enhancer-gene mapping. Many of the predicted kidney function genes are involved in tubule development, emphasizing that developmental effects are likely to be important determinants of adult kidney function.

CRISPRi enhancer-gene mapping identified *NDRG1* and *RBPMS* as kidney function genes. *NDRG1* is expressed during kidney tubule differentiation and was recently demonstrated to participate in hypoxia resistance within the zebrafish pronephros^80,81^. *RBPMS* encodes a regulator of translation and alternative splicing linked to dedifferentiation in multiple cell types^82,83^. It will be interesting to assess whether RBPMS participates in tubule cell dedifferentiation, which in animal models is involved in the development of chronic kidney disease following injury^84^.

Two of the seventeen kidney function cCREs assessed using CRISPRi enhancer-gene mapping affected expression of genes. This proportion is consistent with a prior CRISPRi-based assessment of cCREs^73^. We hypothesize that many cCREs at which perturbation did not identify affected genes in cultured tubule epithelial cells may still regulate tubule gene expression *in vivo*. We suspect that improving the power of CRISPRi screens to identify enhancer-gene pairs will require cellular models that better recapitulate *in vivo* gene regulation, increased sensitivity for detecting changes in genes expressed at moderate and low levels, and more potent CRISPR effectors^85^.

Our approach had several limitations. First, cCREs of cell types or states that were absent or rare in the adult human kidneys would be poorly detected. Undetected cCREs would include those only accessible during kidney development or following injury. These undetected cCREs are likely to account for some of the kidney function heritability not explained by cCREs identified in this study.

Second, while our study uses functionally informed fine-mapping to identify genetic variants likely to influence kidney function, there is no established means of validating which variants are causal. However, several lines of evidence indicate that our fine-mapping identified many of the functional variants: 1) fine-mapped cCRE variants had larger predicted effects onchromatin accessibility than other cCRE variants, 2) fine-mapped variants affected enhancer function, and 3) variants fine-mapped without the use of functional annotation were enriched for missense variants, in kidney tubule cCREs and in conserved regions (Supplementary Figure 5a).

Third, our study focuses on genetic variants associated with kidney function biomarkers used to diagnose kidney disease, but not on kidney disease itself. GWAS of kidney disease have less power than GWAS of kidney function because kidney disease is a binary outcome, in contrast to the continuous variable of eGFR ^6^. Using a GWAS performed in a separate cohort, we found that kidney function variants were associated with chronic kidney disease. This association between kidney function and chronic kidney disease is consistent with prior work showing that loci associated with kidney function biomarkers and chronic kidney disease overlap^6,86^. Thus, the kidney function variants identified in this study are likely to be relevant to chronic kidney disease risk.

## Acknowledgements

G.L. was supported by a T32 from NIDDK and the Physician Scientist Scholars Program from UCSF. This work was also supported by grants from the NIH (R01AR054396 and R01HD089918) to J.F.R. and, in part, by the Laboratory for Genomics Research established by GSK, UCSF and UC Berkeley. We thank Duncan Palmer for graciously sharing code used to generate the Manhattan plot and Nadav Ahituv, David Erle, Katie Pollard, David Morgan, James Pirruccello, Brian Black, and members of the Reiter Laboratory for helpful comments and discussion.

## METHODS

### Bulk ATAC-seq

2g of fresh kidney cortex fragments were homogenized in buffer (250mM sucrose, 10mM Tris pH 7.5, 3mM MgCl2, 0.1% NP40) using a gentleMACS M Tube (Miltenyi Biotec, 130-093-237) and gentleMACS dissociator program E.01. The homogenate was filtered through a 200μm filter, 100μm filter, pelleted by centrifugation, and treated according to the Omni-ATAC protocol^87^, with the following exceptions. Homogenate from 2g of tissue was resuspended in 1ml of ATAC-Resuspension Buffer (RSB) containing detergents. Detergents were diluted with 15ml RSB with 0.1% Tween. Nuclei were passed through a 40μm filter and subsequently counted. 100μl of nuclei were used for the transposition reaction which was performed in 105μl with 10μl of transposase (Illumina, 20034197). The library was sequenced on an Illumina NovaSeq 6000 System on an SP flow cell with paired-end 50-bp reads.

Bulk ATAC-seq alignment to hg38 and peak calling were performed using PEPATAC^88^ (https://pepatac.databio.org/en/latest/). Motif enrichment within ATAC-seq peaks was calculated using HOMER^89^ (http://homer.ucsd.edu/homer/motif/).

### Single-cell ATAC-seq

Libraries were generated with the Chromium Next GEM Single Cell ATAC Library & Gel Bead Kit v1.1 (10x Genomics, 1000176) with the following modifications. Kidney cortex or kidney medulla were dissected and 2 grams of tissue were homogenized and lysed as described for bulk ATAC-seq. Nuclei were either used immediately for scATAC-seq library generation or alternatively 10% DMSO was added to the nuclei suspension and nuclei were frozen in cryovials in a CoolCell Freezing Sytem (Corning) at -80C. To prepare libraries from frozen nuclei, nuclei were thawed at room temperature and immediately diluted 1:8 in diluted nuclei buffer (10x Genomics). To prepare libraries, 200,000 nuclei were pelleted (6 minutes at 500xg, 4C), resuspended in 50uL diluted nuclei buffer, and recounted. 15,000 nuclei were used for transposition and library generation according the manufacturers protocol. Libraries were sequenced on an Illumina NovaSeq 6000 System with paired-end 50-bp reads.

Demultiplexing, read alignment, and cell calling was performed with the Cell Ranger ATAC Pipeline (v1.2, using hg38) to generate scATAC fragment files. Fragment files were loaded into R (3.6.1) using the createArrowFiles function in ArchR (v.0.9.5)^90^. TSS enrichment and number of unique fragments calculated on a per-cell basis were used to remove low quality cells, cutoffs were defined for each library (TSS enrichment >6-7, unique fragments <4000-6800).

Dimensionality reduction, batch effect correction, clustering, integration with kidney scRNA-seq data^26^, and cluster-specific peak calling were performed using the addIterativeLSI, addHarmony, addClusters, adddGeneIntegrationMatrix, and addGroupCoverages functions in ArchR. Gene activity scores were calculated using the ArchR function getMarkerFeatures. Motif enrichment within cell type-specific peaks were identified using the addMotifAnnotations and peakAnnoEnrichment functions and *Homo sapiens* motifs from CIS-BP^91^.

To identify clusters of cell type-specific peaks, the cell type-by-peak matrix was normalized, log transformed, and k-means clustered with 12 clusters. Two clusters corresponded to ubiquitously accessible peaks and were combined.

### Tissue

Deidentified, nontransplantable kidneys were obtained from a not-for-profit organization (DNW, San Ramon, CA, USA). The research protocol was approved by the DNW internal ethics committee (Research project UCSF-18-104). The Institutional Review Board at the University of California San Francisco determined that this project does not meet the definition of human subjects research.

### Chromatin accessibility allelic imbalance

To assess chromatin accessibility allelic imbalance (CAAI), we remapped scATAC reads using WASP^70^, which removes read-mapping bias caused by challenges mapping reads generated from alternate alleles. Sample specific genotypes for using with WASP were generated using Omni 2.5M arrays (Illumina) and imputed using the Michigan Imputation Server^92,93^ (https://imputation.biodatacatalyst.nhlbi.nih.gov/), using TOPMed r2^94^. For each cell type, within each sample reads aligning to heterozygous sites were counted with bcftools mpileup followed by bcftools call. Imputed genotypes were added using bcftools isec. To prevent contamination from incorrectly called heterozygous sites, sites with imputed genotype probability <0.9 and <2 reads from either allele were filtered out. Allelic imbalance significance was assessed with a binomial test, comparing the number of reference reads versus total reads and adjusting for multiple hypothesis testing using the Benjamini-Hochberg method and using an FDR cutoff of 0.01.

To examine transcription factor motifs present at sites with chromatin accessibility allelic imbalance (CAAI), FIMO was used to scan the 20 bp region surrounding each variant for matches in the JASPAR 2020 CORE non-redundant vertebrates database^95,96^. For each variant, we queried both the reference and alternate sequence, keeping matches with a p-value of less than 1e-3 for either the reference or alternate sequence. To identify motifs affected by variants, only matches satisfying 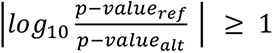 were analyzed. Enrichment of motifs affected by variants at CAAI variants was determined by comparing against variants without CAAI using a hypergeometric test. P-values were corrected for multiple hypothesis testing using the Benjamini-Hochberg procedure. To examine the direction of effect of motifs on chromatin accessibility, for each motif predicted to be affected at sites with CAAI, we determined the fraction of sites where the motif is predicted by FIMO to preferentially bind to the allele associated with higher chromatin accessibility.

Only transcription factor motifs that correspond to one of the top 300 TFs expressed in the indicated cell type were plotted^26^.

### Partitioned Heritability Analyses

Partitioned heritability analyses were performed using stratified LD score regression^22^ using baseline annotations (baselineLD_v2.2, https://alkesgroup.broadinstitute.org/LDSCORE/)^97^.

### GWAS

The GWAS of eGFR_cr-cys_ in the UK Biobank^98^ (release version 3) was performed in unrelated individuals of European ancestry using the inverse-rank normal transformation of eGFR_cr-cys_ defined using the 2012 CKD-EPI equation^14^. Age, sex, and the fist 10 principal components were used as covariates.

To identify eGFR_cr-cys_ loci, the variant with the lowest *P* value across the genome was defined as an index variant and a 1 Mb locus centered at the index variant was defined. This was repeated until no further variants with *P* values < 5×10^−8^ remained. Each locus was then classified in one of four categories to indicate whether it contained a genome-wide significant variant for the eGFR_cr_ and eGFR_cys_ phenotype: 1) contains a genome-wide significant variant for eGFR_cr_ and eGFR_cys_, 2) contains a genome-wide significant variant for eGFRcr, 3) contains a genome-wide significant variant for eGFR_cys_, 4) locus unique to eGFR_cr-cys_. Overlapping loci within the same category were merged. The small number of overlapping loci within different categories were trimmed at the midpoint of the overlap.

### Gene prioritization

We applied PoPS^60^ to the eGFR_cr-cys_ GWAS to prioritize kidney function genes. We calculated per-gene association statistics using MAGMA^99^ with the 1000G EUR reference panel^100^ and Ensembl 107 gene coordinates. We then learned a gene prioritization score using PoPS from the MAGMA statistics and 57,543 PoPS gene features (https://www.finucanelab.org/data). To prioritize genes, we selected the top PoPS scoring gene within 500kb of a fine-mapped variant (PoPS score), the nearest gene body to the fine-mapped variant (distance), or only selected the gene if the top PoPS scoring gene and closest gene were the same (PoPSxDistance).

Gene prioritization was evaluated with two sets of gold-standard kidney function genes 1) genes with fine-mapped missense variants with PIP>0.7 from the eGFR_cr-cys_ GWAS (an approach analogous to the one used in the original PoPS manuscript) and 2) genes with coding mutations associated with both creatinine and cystatin C levels from the UK Biobank exome data assessed by gene-based association testing^101^. To define the exome based set we used the gene-based association (SKAT-O test) between putative loss of function variants and both creatinine and cystatin C levels (https://genebass.org). To exclude genes associated only with either creatinine or cystatin C, we used the maximum(p-value creatinine, p-value cystatin C). These p-values were adjusted for multiple hypothesis testing (Benjamini-Hochberg). This procedure was repeated for the gene-based association for missense variants. Genes with FDR<0.1 from either the putative loss of function or missense tests were included in the gold standard set.

To evaluate gene prioritization approaches, we defined test loci as the 500kb region around each fine-mapped noncoding variant (PIP>0.4) that overlapped a gold-standard gene. If the locus for two fine-mapped variants overlapped, one was chosen randomly to be part of the test set. For each gold standard and gene prioritization metric, we quantified the precision and recall. Precision was defined as the proportion of prioritized genes that are in the gold standard set. Recall was defined as the number of gold standard genes in the tested loci that were prioritized.

### Functionally informed fine-mapping

eGFR_cr-cys_ loci for fine-mapping were identified by identifying by defining loci containing an index variant associated with both creatinine and cystatin C. Index eGFR_cr-cys_ variants from the eGFR_cr-cys_ GWAS were identified using GCTA-COJO. Association of these index eGFR_cr-cys_ variants with both creatinine and cystatin C was assessed using summary statistics from the GWAS for the individual biomarkers performed in UK Biobank. Fine-mapping was performed on 1 Mb loci centered on index eGFR_cr-cys_ variants with nominally significant p-values for both individual biomarkers (Bonferroni corrected for the 522 tests, p<0.05) and the same direction of effects for both biomarkers. Fine-mapping of these loci was performed using Polyfun^38^ and Susie^102^ using a maximum of 3 causal variants per locus. Standard fine-mapping was performed using Susie, functionally informed fine-mapping used prior causal probabilities calculated in Polyfun based on both a kidney epithelial regulatory element annotation and baseline annotations^38^.

### Deep learning prediction of chromatin accessibility from DNA sequence (ChromKid)

We trained a set of multi-task convolutional neural networks (CNN), which we collectively refer to as ChromKid, to map 1344 bp DNA sequence regions across the genome to quantitative read outs of chromatin accessibility across 10 kidney cell types. For efficient training, we used a transfer learning procedure (see below) where we first trained a classification model to predict binary labels (accessible vs. not accessible) and then fine-tuned a regression model to predict quantitative read outs. We trained 23 CNNs using a leave-one-chromosome-out (LOCO) approach, described below, to prevent overfitting.

Model architecture: We used the Basenji repository (https://github.com/calico/basenji) and a variation of the previously optimized multi-task Basset CNN architecture for predicting genome-wide chromatin accessibility from DNA sequence. The inputs to the model are 1344 bp long DNA sequences that are one-hot encoded. The model is composed of 8 convolutional layers followed by 2 fully connected layers.

The first convolutional layer has 288 filters with a kernel size of 17, and is followed by max pooling with size 3. The six subsequent convolutional layers each have consecutively increasing numbers of filters, beginning with 288 and ending with 512. The increasing number of filters in the convolutional layers are defined as 1.122 times the number of filters in the previous layer. Each of these six convolutional layers have a kernel size of 5 and are followed by max pooling with size 2. The final convolutional layer has 256 filters with a kernel size of 1. Each convolutional layer is followed by batch normalization and a GELU non-linearity.

The convolutional layers are followed by a fully connected layer with 768 neurons. This layer is followed by batch normalization and a GELU non-linearity, and has a dropout probability of 0.2.

The final fully connected layer is the output layer. This layer maps to 10 outputs (multi-task output), each corresponding to a read out of chromatin accessibility in a particular cell type. We use binary or continuous output labels and associated loss functions in the multi-stage training (see below). When training on binary labels (accessible vs. not accessible), we use the binary cross-entropy loss function with sigmoid outputs. When training on continuous, quantitative measures of accessibility, we use the poisson loss function with softplus outputs.

Data processing: To train and evaluate the classification model, ATAC-seq peak calls were used. Genomic contigs containing a peak were binned into 1344 bp windows with a stride of 192 bp. Binary labels were assigned corresponding to whether the central 192 bp of each window overlap an ATAC-seq peak by at least 25%. To train and evaluate the regression model, continuous, quantitative ATAC-seq values were used. Continuous valued labels were assigned by summing the read counts in the central 192 bp of each window. In both cases, genome assembly gaps and regions of the genome previously blacklisted owing to signal artifact are removed^103^.

Training & transfer learning procedure: We trained 23 CNNs with the architecture described above, using a LOCO approach to prevent overfitting. For each of the 23 CNNs, one chromosome was completely held out during training and validation. The X and Y chromosomes were grouped together and both held out from the same model. For all models where an odd numbered chromosome was held out, and for the model where the X and Y chromosomes were held out, chromosomes 2, 18, and 22 were used for validation. For all models where an even numbered chromosome was held out, chromosomes 5, 9, and 21 were used for validation. For each model, the same training, validation, and test split was used for both training stages to ensure that the test set was completely held-out throughout training.

For each of the 23 CNNs, we first train a multi-task classification model with the architecture and data described above. The models were trained using the Keras SGD optimizer (learning rate = 0.005), binary cross-entropy loss, and early stopping with patience of 3 epochs. We then fine-tune a set of multi-task regression models with the architecture described above. We begin by initializing each model with the parameters learned by the corresponding classification model. We “freeze” the parameters of the convolutional layers, such that parameters in the frozen layers cannot be altered during subsequent training. We fine-tune the parameters of the fully connected layers by training on the continuous valued data described above. The models were trained using the Keras SGD optimizer (learning rate = 0.005), Poisson loss, and early stopping with patience of 3 epochs. All models were trained on Nvidia 1080ti GPUs.

To evaluate the genome-wide performance of the models, we treat all genomic sequences processed using the procedure described above as our test set. For each sequence, we predict quantitative read outs of chromatin accessibility using the model where the relevant chromosome was held out from training.

Prediction of chromatin accessibility allelic imbalance: To quantify ChromKid variant effect performance, we identified variants with allelic imbalance in each of proximal tubule, loop of Henle, and distal tubule as described above. Non-allelic imbalance sets were constructed using variants where the p-value was greater than 0.1. The non-allelic imbalance sets were adjusted to be as large as possible while matching the read count distribution of the positive set variants on a log scale, resulting in a non-allelic imbalance set that was 7x the size of the allelic imbalance set in each of LOH, PT, and DT.

ChromKid predictions for the reference and alternate alleles in the 192 bp bin centered at the variant were defined as REF and ALT, and the predicted CAAI was computed as REF / (REF + ALT). We evaluated how well predicted CAAI could classify whether a variant had CAAI (i.e., discriminate CAAI variants from non-CAAI variants) using the area under the receiver operating characteristic curve (AUROC). As an additional metric, among the variants in the CAAI set, we computed the AUROC for prediction of CAAI direction (REF>ALT vs ALT>REF) as measured by AUROC.

### SNP Activity Difference Tracks

To quantify the predicted change in chromatin accessibility due to a single nucleotide variant (SNV), we calculated the SNV activity difference (SAD) score as the difference between the predicted ATAC-seq coverage in a 192-bp bin for the alternate and reference alleles^103^. To compute a SAD score track for a given SNV, we plotted the model’s predicted SAD score centered at every position for which the SNV falls within the model’s receptive field. This resulted in predictions for each cell type at 1344 positions, representing the model’s 1344-bp receptive field.

### Enhancer Assays

Regulatory elements were amplified by PCR from human genomic DNA using primers listed in Supplementary Table 10 and cloned into the pGL4.23 Luciferase Reporter Vector (Promega, E8411) linearized with NheI and EcoRV (New England Biolabs) using In-Fusion Snap Assembly (Takara, 638949). To obtain regulatory elements with variant alleles not present in the genomic DNA used for amplification, site-directed mutagenesis was performed using In-Fusion Snap Assembly using primers listed in Supplementary Table 10. Sequences of all constructs were verified using long read sequencing (Primordium Labs), which confirmed that reference and alternate allele plasmids only different at the specified variant. Plasmids containing regulatory elements were transfected into Human Renal Cortical Epithelial cells (HRCE; Lonza, CC-2554). Specifically, 1.25 × 10^4^ HRCE cells were plated in a 96-well plate in renal epithelial growth media (Lonza, CC-3190). 24 hours later cells were transfected with 50ng regulatory element plasmid and 50ng Renilla plasmid (pGL 4.74; Promega, E6921) using 0.3µL Fugene HD (Promega, E2311). 24h post-transfection, regulatory element activity was measured using the Dual-Luciferase Reporter Assay (Promega, E1910) and a GloMax 96 Luminometer according to manufacturer’s instructions. Regulatory element activity was plotted as the ratio of firefly to luciferase signal.

### CRISPRi enhancer-gene mapping

sgRNAs were cloned into a BFP expressing dual guide expression vector, a modified version of PLGR002 which contains a second guide cassette (Addgene, 188320). Both guides within a vector were designed to target either: 1) A cCRE harboring a fine-mapped kidney function variant (17 cCREs, 2 plasmids per cCRE), 2) the transcription start site of a gene (6 TSS controls), 3) a known enhancer^73^ that we had previously validated in HRCEs (7 control enhancers, 2 plasmids per enhancer), or 4) a negative controls: gene desert, closed chromatin, or non-targeting (12 negative controls). All guides were chosen using CRISPick^104^ with the exception of TSS guides which were chosen from a published guide library^105^. Lentivirus was produced from sgRNA plasmids by co-transfecting Lenti-X 293T (Takarabio, 632180) cells with the sgRNA plasmid library and packaging plasmids-psPAX2 and pMD2.G (Addgene, 12260 and 12259) using Trans-IT-293 transfection reagent (Mirus, MIR2700). Media was collected on day 2 after transfection and the lentivirus was concentrated using Lenti-X Concentrator (Takarabio, 631232) and resuspended in renal epithelial growth media. The same protocol was used to generate lentivirus from the CBH-dCas9-mCherry-ZIM3 plasmid, which was generated using the ZIM3 sequence from pLX303-ZIM3-KRAB-dCAS9 (Addgene, 154472)^106^.

HRCEs (Lonza, CC-2554) from 4 donors grown in renal epithelial growth media (Lonza, CC-3190) were transduced with lentivirus encoding dCas9-mCherry-ZIM3. Three days later cells were transduced with lentivirus encoding the sgRNA library and blue fluorescent protein (BFP). Nine days after guide transduction mCherry, BFP double-positive cells from each donor were sorted. Equal numbers of double-positive cells from each donor were pooled and prepared for single cell RNA-sequencing, with 4 GEM groups each loaded with 20,000 cells, using the Chromium Next GEM Single Cell 3’ Kit 3.1 (10x Genomics, 1000268) according to the 10x Genomics protocol for Feature Barcode technology for CRISPR Screening. The resulting guide and expression libraries were sequenced on a Novaseq 6000 (Illumina).

Read alignment and cell assignment was performed using Cellranger count v7.0.0, the GRCh38-2020-A human reference transcriptome from 10x, and the protospacer sequences (Supplementary Table 9)^107^. Cell doublets were identified using Souporcell using the number of clusters argument -k 4^108^. Doublets were discarded.

Seurat v4.3.0 was used for subsequent analysis^109^. Cells with mitochondrial RNA percentage, RNA count, or number of detected genes outside three median absolute deviations were filtered out. Seurat SCTransform v2 was used to normalize and scale the data as well as to find the variable features. The top 3000 highly variable genes were used for PCA and UMAP dimensionality reductions. Libraries from each screen were integrated using Seurat’s SelectIntegrationFeatures, FindIntegrationAnchors, and IntegrateData functions. Cells were clustered using FindClusters function and the Louvain algorithm.

Guide calling was performed using a Poisson-Gaussian caller^110,111^. Differential expression testing was performed using Seurat’s FindMarkers function and Logistic Regression test with library and donor as latent variables, and log fold change threshold of 0.03. Only genes within 1 Mbase up/downstream of the target (distal element or TSS) were considered for the test. Reported p-values were Bonferroni-corrected by the total number of neighbors of all targets in the experiment. MAST and Wilcoxon Rank Sum tests yielded similar results.

## Data Availability Statement

Kidney ATAC-seq and scATAC-seq data is deposited with the Gene Expression Omnibus (GEO) (accession no. GSE262931).

## Code Availability Statement

Code used for the analysis of scATAC-seq data, allelic imbalance, prediction of chromatin accessibility and variant effect from sequence (ChromKid), and benchmarking PoPS for kidney function are available at GitHub repository (https://github.com/ni-lab/kidney-finemapping). Code used for CRISPRi ehancer-gene mapping are available at GitHub repository (https://github.com/ucsf-lgr/ckd-workflow).

---

**Supplementary Figure 1:**
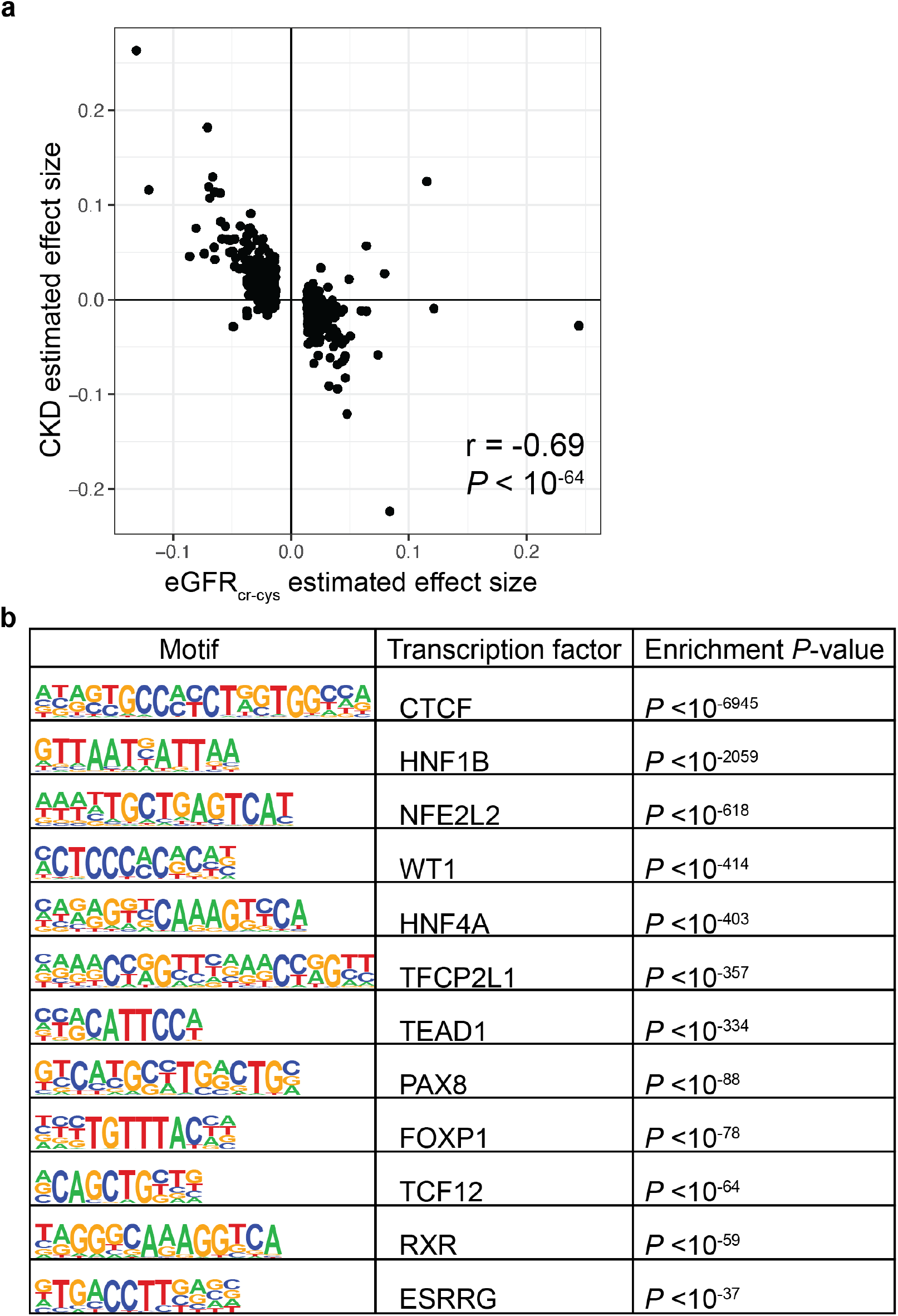
Kidney candidate *cis-*regulatory elements are enriched for kidney function heritability. **a)** Pearson correlation between variant effect size for eGFR_cr-cys_ and chronic kidney disease (CKD) from GWAS performed in two independent populations. The *P* value for the Pearson correlation is indicated. **b)** Binding motifs of kidney transcription factors are enriched in kidney cCREs defined using kidney ATAC-seq peaks. *P* values are calculated relative to GC-content matched genomic background using the binomial test.

**Supplementary Figure 2:**
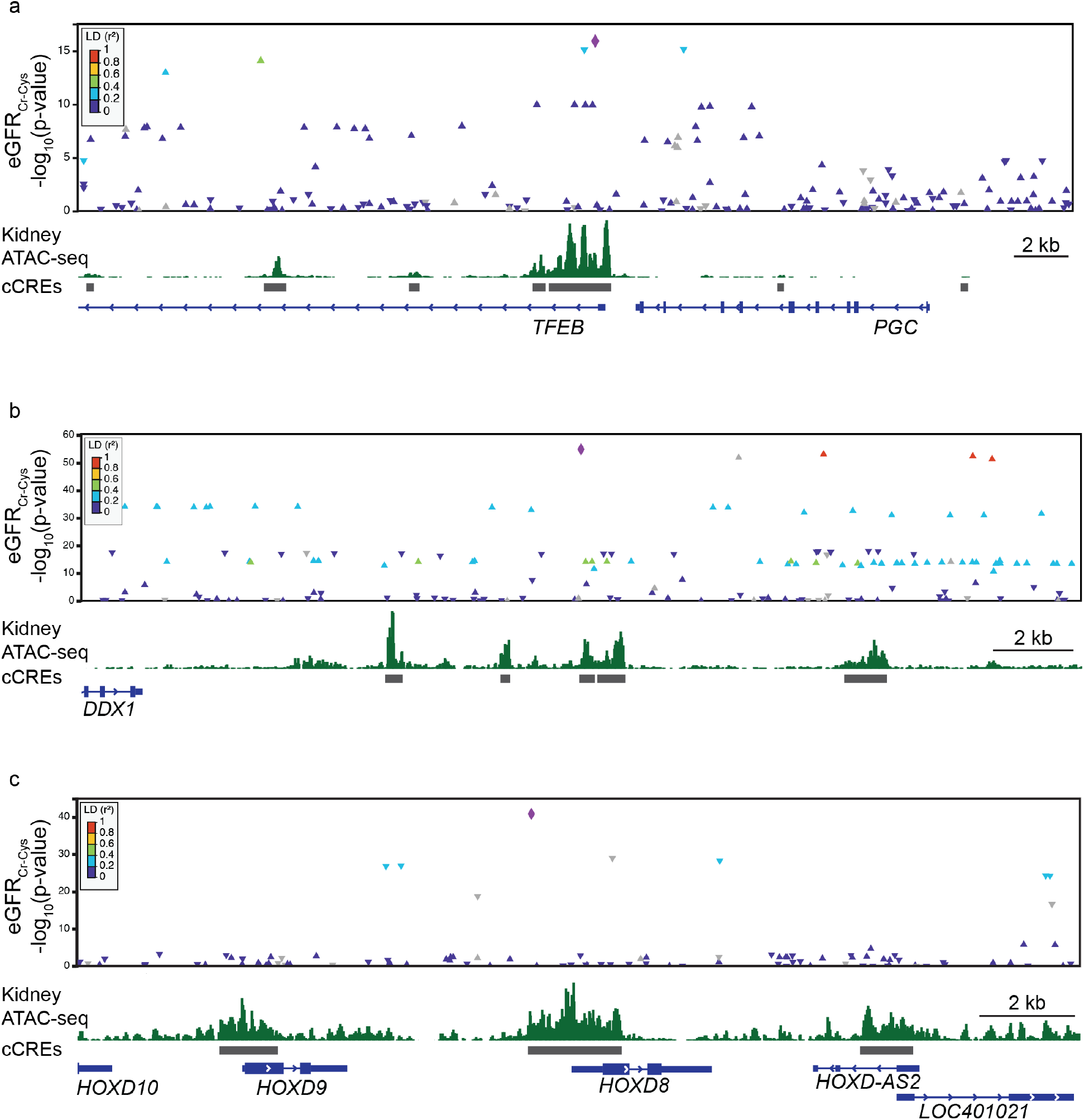
Kidney function variants are within kidney candidate *cis*-regulatory elements. **a-c)** The association between eGFR_cr-cys_ and variants near *TFEB*, *DDX1* and *HOXD8* are indicated. The variant with the strongest association at each locus is indicated by the purple diamond and the LD between the top variant and other variants at the locus are indicated by color.

**Supplementary Figure 3:**
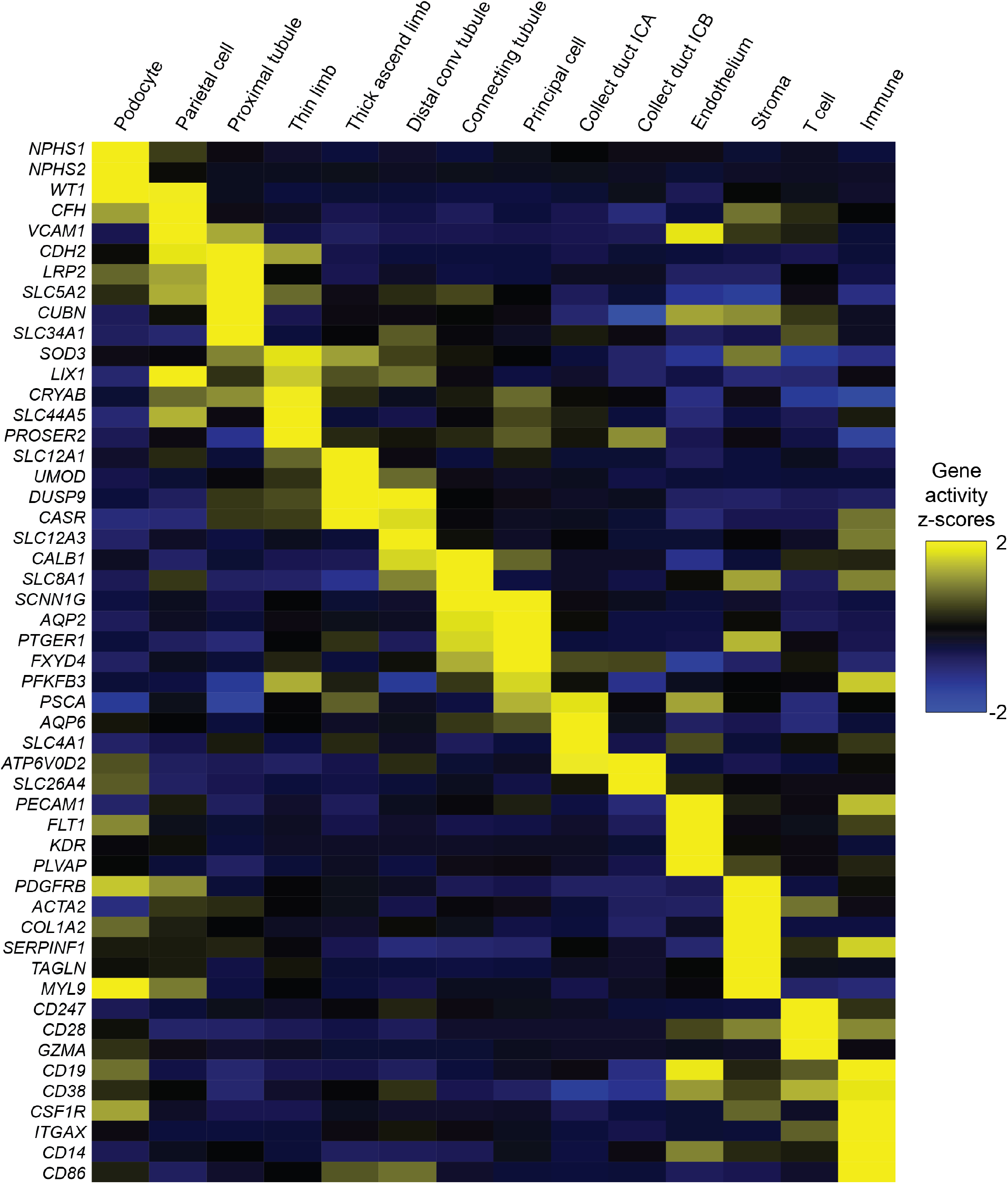
Gene activity scores of kidney marker genes. Gene activity scores—a measure of integrated chromatin accessibility around genes which correlates with expression—for established marker genes of each cell type.

**Supplementary Figure 4:**
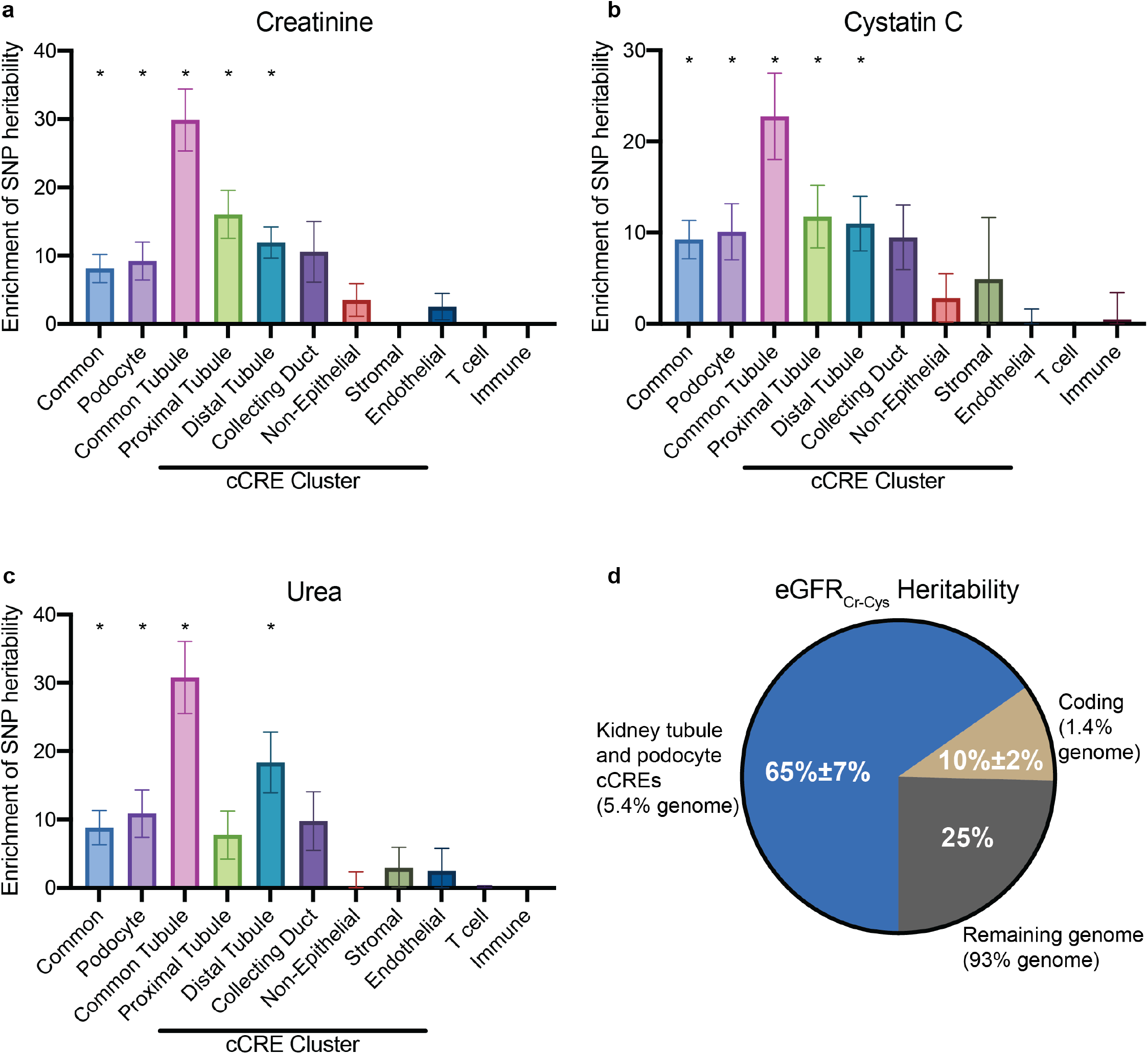
Kidney tubule epithelial cCREs account for most kidney function SNP heritability. **a-c)** Fold enrichment of SNP heritability for serum creatinine, cystatin C, and urea levels within cell type-specific and ubiquitously accessible (common) cCREs calculated using stratified LD score regression. * *P*<0.05. *P* values are Bonferroni corrected for multiple hypothesis testing. **d)** Fraction of heritability of eGFR_cr-cys_ explained by kidney podocyte and tubule epithelial cell cCREs (blue), coding exons (beige), and the remainder of the human genome (gray).

**Supplementary Figure 5:**
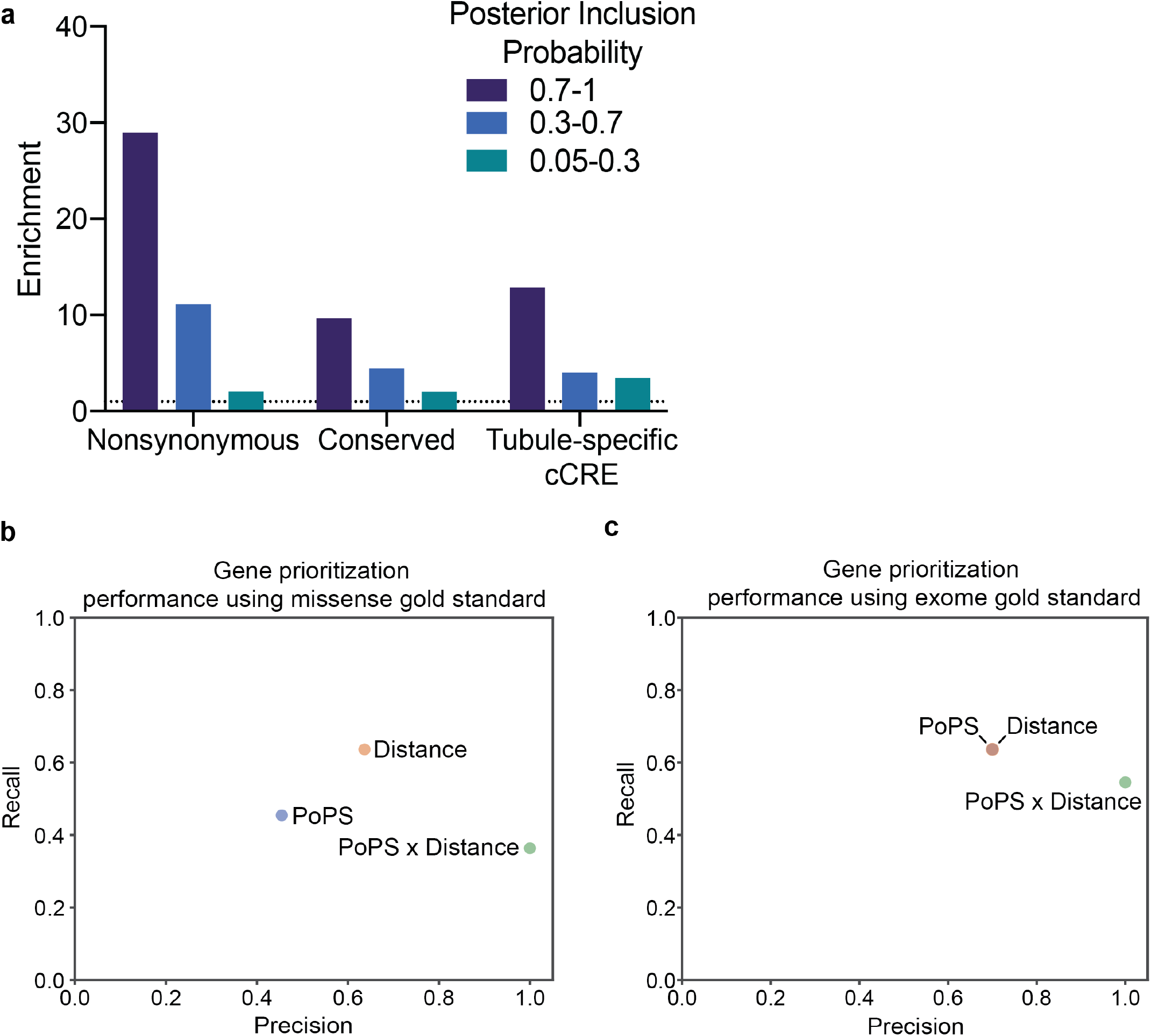
Functionally informed fine-mapping and target gene prioritization identifies causal kidney function variants and genes. **a)** Enrichment of variants fine-mapped without functional annotations. Fold enrichment for variants that cause nonsynonymous changes in a coding sequence, are evolutionarily conserved, or that lie within tubule epithelial-specific cCREs is shown. Variants are separated into those with high (>0.7), moderate (0.3-0.7), and low (0.05-0.3) posterior inclusion probability. **b-c)** Precision and recall of the top prioritized gene for fine-mapped variants (PIP>0.4) based on distance, PoPS, or the agreement of PoPS and distance using a gold standard defined as (b) genes with fine-mapped missense variants associated with eGFR_cr-cys_ or (c) genes significantly associated with both creatinine and cystatin C in an exome sequencing gene-based association test.

**Supplementary Figure 6:**
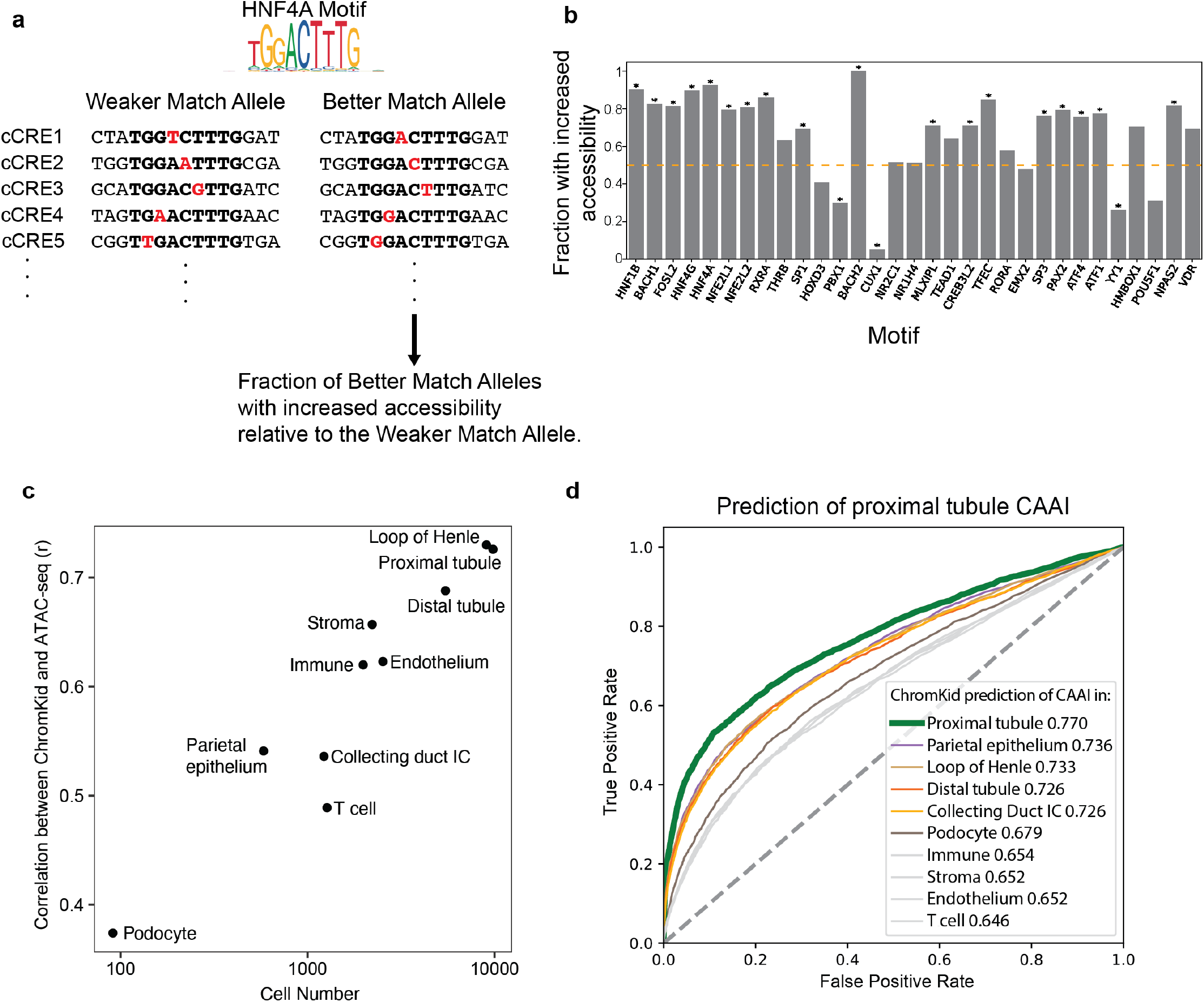
Chromatin accessibility allelic imbalance and machine learning identify cell type-specific effects of variants on chromatin accessibility. **a)** Schematic illustrating alleles that differ in their similiarity to the HNF4A transcription factor binding motif. **b)** The fraction of cCREs at which the variant more closely matching the consensus binding motif of the indicated transcription factor is associated with increased chromatin accessibility. **c)** The model Pearson correlation between ChromKid-predicted and experimentally measured chromatin accessibility calculated on a held-out chromosome (chromosome 11) graphed relative to the number of cells within the training set for each cell type. Across cell types, correlation increased with greater cell number. **d)** ROC curves for the prediction of proximal tubule chromatin accessibility (CAAI), displaying the false positive rate (x-axis) versus true positive rate (y-axis) of cell-type specific predictions from ChromKid. The AUCs for ChromKid predictions for each cell type are shown in the legend.

**Supplementary Figure 7:**
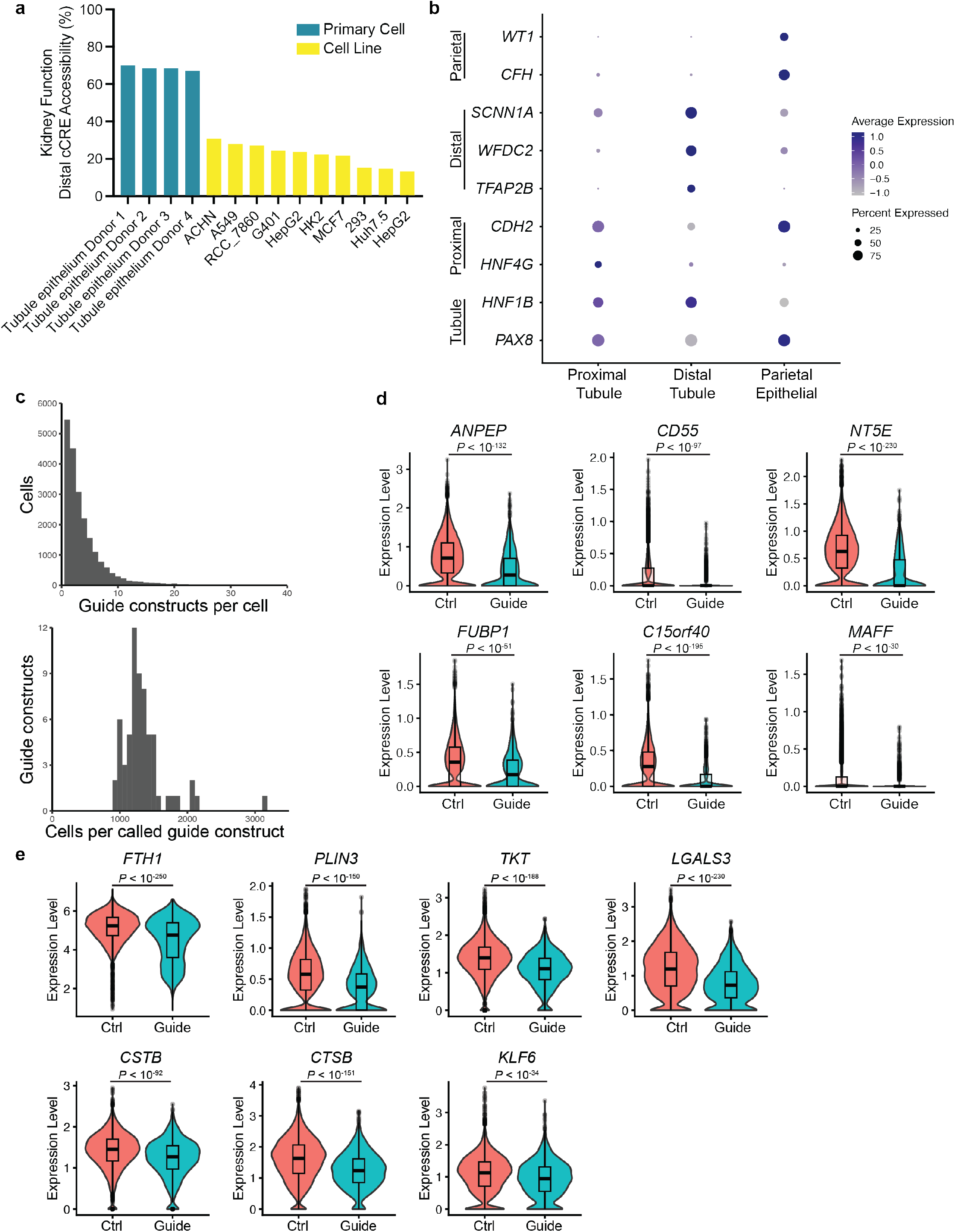
CRISPRi enhancer-gene mapping in human tubule epithelial cells. **a)** The fraction of non-promoter tubule epithelial cCREs containing fine-mapped kidney function variants (PIP>0.4) which are accessible in cultured primary human tubule epithelial cells and indicated cell lines. **b)** Expression of marker genes in proximal tubule, distal tubule, or parietal epithelial cells within cultured primary human tubule epithelial cells as measured by scRNA-seq. Marker genes of tubule cells, proximal tubule, distal tubule, and parietal epithelial cells are indicated on the y-axis. **c)** Histograms of the number of guide constructs expressed per cell (top) and number of cells expressing each guide construct (bottom). **d)** Effect of transcriptional start site-targeting guides on gene expression. Expression of indicated genes in cells that do not express (Ctrl) or do express a guide (Guide) targeting the transcription start site. **e)** Effect of positive control enhancer guides on gene expression. Expression of indicated genes in cells that do not express (Ctrl) or do express a guide (Guide) targeting a previously identified enhancer. Superimposed box plots indicate the median, 25^th^ and 75^th^ percentiles of expression. All *P* values were calculated using logistic regression and adjusted for multiple hypothesis testing using the Bonferroni correction.

**Supplementary Figure 8:**
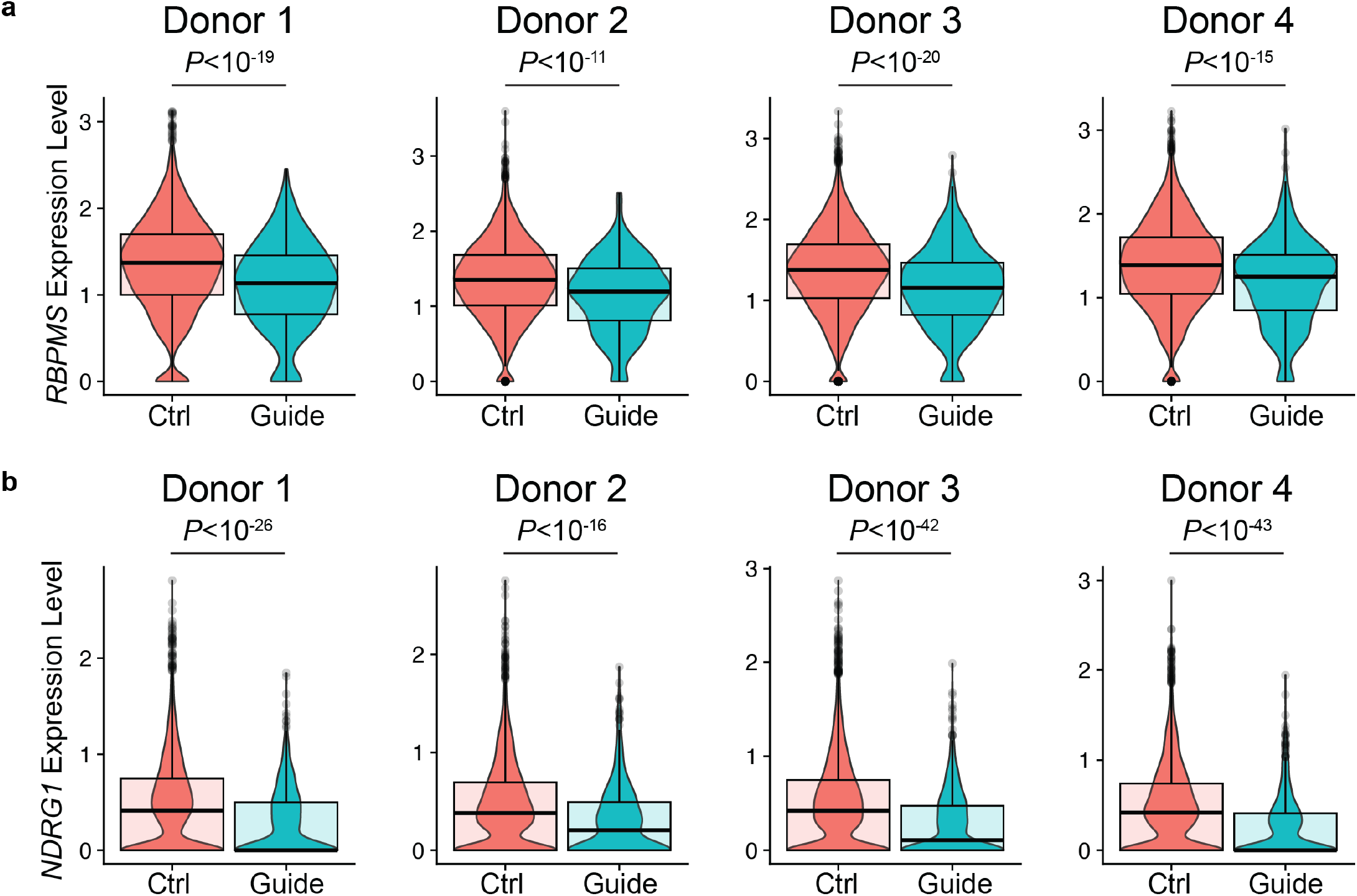
Consistency of the effect of cCRE silencing across donors. **a)** Violin plots of *RBPMS* gene expression in cells expressing (Guide) or not expressing (Ctrl) guides targeting the cCRE within the first intron of *RBPMS* for four different donors. **b)** Violin plots of *NDRG1* gene expression in cells expressing (Guide) or not expressing (Ctrl) guides targeting the cCRE 5’ of *NDRG1* in HRCEs. Superimposed box plots indicate the median, 25^th^ and 75^th^ percentiles of expression. All *P* values were calculated using the logistic regression test and adjusted for multiple hypothesis testing using the Bonferroni correction.

## Notes

### Competing Interest Statement

Volkan Sevim, Audrey Chu, Jonathan Davitte, and Radu Rapiteanu are employees of GlaxoSmithKline

## References

1. Dantzler, W. H. Comparative Physiology of the Vertebrate Kidney. (Springer Science & Business Media, 2012).

2. Cockwell, P. & Fisher, L.-A. The global burden of chronic kidney disease. The Lancet 395, 662–664 (2020).

3. Akrawi, D. S. et al. Heritability of End-Stage Renal Disease: A Swedish Adoption Study. Nephron 138, 157–165 (2018).

4. Arpegård, J. et al. Comparison of heritability of Cystatin C- and creatinine-based estimates of kidney function and their relation to heritability of cardiovascular disease. J. Am. Heart Assoc. 4, e001467 (2015).

5. Raggi, P. et al. Heritability of renal function and inflammatory markers in adult male twins. Am. J. Nephrol. 32, 317–323 (2010).

6. Wuttke, M. et al. A catalog of genetic loci associated with kidney function from analyses of a million individuals. Nat. Genet. 51, 957–972 (2019).

7. Hellwege, J. N. et al. Mapping eGFR loci to the renal transcriptome and phenome in the VA Million Veteran Program. Nat. Commun. 10, 3842 (2019).

8. Pattaro, C. et al. Genetic associations at 53 loci highlight cell types and biological pathways relevant for kidney function. Nat. Commun. 7, 10023 (2016).

9. Köttgen, A. et al. Multiple loci associated with indices of renal function and chronic kidney disease. Nat. Genet. 41, 712–717 (2009).

10. Stevens, L. A., Coresh, J., Greene, T. & Levey, A. S. Assessing Kidney Function — Measured and Estimated Glomerular Filtration Rate. N. Engl. J. Med. 354, 2473–2483 (2006).

11. Sieber, K. B. et al. Integrated Functional Genomic Analysis Enables Annotation of Kidney Genome-Wide Association Study Loci. J. Am. Soc. Nephrol. 30, 421 (2019).

12. Liu, H. et al. Epigenomic and transcriptomic analyses define core cell types, genes and targetable mechanisms for kidney disease. Nat. Genet. 54, 950–962 (2022).

13. Sheng, X. et al. Mapping the genetic architecture of human traits to cell types in the kidney identifies mechanisms of disease and potential treatments. Nat. Genet. 53, 1322–1333 (2021).

14. Inker, L. A. et al. Estimating Glomerular Filtration Rate from Serum Creatinine and Cystatin C. N. Engl. J. Med. 367, 20–29 (2012).

15. Inker, L. A. et al. New Creatinine- and Cystatin C–Based Equations to Estimate GFR without Race. N. Engl. J. Med. 385, 1737–1749 (2021).

16. Bekheirnia, M. R. et al. Whole-exome sequencing in the molecular diagnosis of individuals with congenital anomalies of the kidney and urinary tract and identification of a new causative gene. Genet. Med. Off. J. Am. Coll. Med. Genet. 19, 412–420 (2017).

17. Niranjan, T. et al. The Notch pathway in podocytes plays a role in the development of glomerular disease. Nat. Med. 14, 290–298 (2008).

18. Bonegio, R. G. B., Beck, L. H., Kahlon, R. K., Lu, W. & Salant, D. J. The fate of Notch-deficient nephrogenic progenitor cells during metanephric kidney development. Kidney Int. 79, 1099–1112 (2011).

19. Zewinger, S. et al. Apolipoprotein C3 induces inflammation and organ damage by alternative inflammasome activation. Nat. Immunol. 21, 30–41 (2020).

20. Carew, R. M. et al. Deletion of Irs2 causes reduced kidney size in mice: role for inhibition of GSK3β? BMC Dev. Biol. 10, 73 (2010).

21. Kakoki, M., McGarrah, R. W., Kim, H.-S. & Smithies, O. Bradykinin B1 and B2 receptors both have protective roles in renal ischemia/reperfusion injury. Proc. Natl. Acad. Sci. 104, 7576–7581 (2007).

22. Finucane, H. K. et al. Partitioning heritability by functional annotation using genome-wide association summary statistics. Nat. Genet. 47, 1228–1235 (2015).

23. Napolitano, G. et al. A substrate-specific mTORC1 pathway underlies Birt–Hogg–Dubé syndrome. Nature 585, 597–602 (2020).

24. Di-Poï, N., Zákány, J. & Duboule, D. Distinct Roles and Regulations for Hoxd Genes in Metanephric Kidney Development. PLOS Genet. 3, e232 (2007).

25. Granja, J. M. et al. ArchR is a scalable software package for integrative single-cell chromatin accessibility analysis. Nat. Genet. 53, 403–411 (2021).

26. Wilson, P. C. et al. The single-cell transcriptomic landscape of early human diabetic nephropathy. Proc. Natl. Acad. Sci. 116, 19619–19625 (2019).

27. Marable, S. S., Chung, E., Adam, M., Potter, S. S. & Park, J.-S. Hnf4a deletion in the mouse kidney phenocopies Fanconi renotubular syndrome. JCI Insight 3, e97497.

28. Chen, L. et al. Transcriptomes of major renal collecting duct cell types in mouse identified by single-cell RNA-seq. Proc. Natl. Acad. Sci. 114, (2017).

29. Scott, E. W., Simon, M. C., Anastasi, J. & Singh, H. Requirement of transcription factor PU.1 in the development of multiple hematopoietic lineages. Science 265, 1573–1577 (1994).

30. Somlo, S. & Mundel, P. Getting a foothold in nephrotic syndrome. Nat. Genet. 24, 333–335 (2000).

31. Gekle, M. Renal Tubule Albumin Transport. Annu. Rev. Physiol. 67, 573–594 (2005).

32. Reginensi, A. et al. SOX9 controls epithelial branching by activating RET effector genes during kidney development. Hum. Mol. Genet. 20, 1143–1153 (2011).

33. Hart, T. C. et al. Mutations of the UMOD gene are responsible for medullary cystic kidney disease 2 and familial juvenile hyperuricaemic nephropathy. J. Med. Genet. 39, 882–892 (2002).

34. Kang, H. M. et al. Sox9-Positive Progenitor Cells Play a Key Role in Renal Tubule Epithelial Regeneration in Mice. Cell Rep. 14, 861–871 (2016).

35. Wyss, M. & Kaddurah-Daouk, R. Creatine and Creatinine Metabolism. Physiol. Rev. 80, 1107–1213 (2000).

36. Lepist, E.-I. et al. Contribution of the organic anion transporter OAT2 to the renal active tubular secretion of creatinine and mechanism for serum creatinine elevations caused by cobicistat. Kidney Int. 86, 350–357 (2014).

37. Tanihara, Y. et al. Substrate specificity of MATE1 and MATE2-K, human multidrug and toxin extrusions/H(+)-organic cation antiporters. Biochem. Pharmacol. 74, 359–371 (2007).

38. Weissbrod, O. et al. Functionally informed fine-mapping and polygenic localization of complex trait heritability. Nat. Genet. 52, 1355–1363 (2020).

39. Li, M., Li, Y., Liu, Y., Zhou, X. & Zhang, H. An Updated Review and Meta Analysis of Lipoprotein Glomerulopathy. Front. Med. 9, (2022).

40. Ali, M., Rellos, P. & Cox, T. M. Hereditary fructose intolerance. J. Med. Genet. 35, 353–365 (1998).

41. Clissold, R. L., Hamilton, A. J., Hattersley, A. T., Ellard, S. & Bingham, C. HNF1B-associated renal and extra-renal disease—an expanding clinical spectrum. Nat. Rev. Nephrol. 11, 102–112 (2015).

42. Figueres, M.-L. et al. Kidney Function and Influence of Sunlight Exposure in Patients With Impaired 24-Hydroxylation of Vitamin D Due to CYP24A1 Mutations. Am. J. Kidney Dis. 65, 122–126 (2015).

43. Kantarci, S. et al. Mutations in LRP2, which encodes the multiligand receptor megalin, cause Donnai-Barrow and facio-oculo-acoustico-renal syndromes. Nat. Genet. 39, 957–959 (2007).

44. Dasgupta, D. et al. Mutations in SLC34A3/NPT2c Are Associated with Kidney Stones and Nephrocalcinosis. J. Am. Soc. Nephrol. 25, 2366 (2014).

45. Karet, F. E. et al. Mutations in the gene encoding B1 subunit of H+-ATPase cause renal tubular acidosis with sensorineural deafness. Nat. Genet. 21, 84–90 (1999).

46. Lemaire, M. Novel Fanconi renotubular syndromes provide insights in proximal tubule pathophysiology. Am. J. Physiol.-Ren. Physiol. 320, F145–F160 (2021).

47. Ward, C. J. et al. The gene mutated in autosomal recessive polycystic kidney disease encodes a large, receptor-like protein. Nat. Genet. 30, 259–269 (2002).

48. The European Polycystic Kidney Disease Consortium. The polycystic kidney disease 1 gene encodes a 14 kb transcript and lies within a duplicated region on chromosome 16. Cell 77, 881–894 (1994).

49. Feliubadaló, L. et al. Non-type I cystinuria caused by mutations in SLC7A9, encoding a subunit (bo,+AT) of rBAT. Nat. Genet. 23, 52–57 (1999).

50. Sun, Z. et al. A genetic screen in zebrafish identifies cilia genes as a principal cause of cystic kidney. Development 131, 4085–4093 (2004).

51. Messaoudi, S. et al. Endothelial Gata5 transcription factor regulates blood pressure. Nat. Commun. 6, 8835 (2015).

52. Wang, C. et al. Loss of DEPTOR in renal tubules protects against cisplatin-induced acute kidney injury. Cell Death Dis. 9, 1–11 (2018).

53. Centini, R. et al. Loss of Fnip1 alters kidney developmental transcriptional program and synergizes with TSC1 loss to promote mTORC1 activation and renal cyst formation. PLoS ONE 13, e0197973 (2018).

54. López-Rodríguez, C. et al. Loss of NFAT5 results in renal atrophy and lack of tonicity-responsive gene expression. Proc. Natl. Acad. Sci. 101, 2392–2397 (2004).

55. Schell, C., et al. The FERM protein EPB41L5 regulates actomyosin contractility and focal adhesion formation to maintain the kidney filtration barrier. Proc. Natl. Acad. Sci. 114, E4621–E4630 (2017).

56. Zhang, J. et al. TWIK-related acid-sensitive K+ channel 2 promotes renal fibrosis by inducing cell-cycle arrest. iScience 25, 105620 (2022).

57. Ma, M. K. M., Yung, S. & Chan, T. M. mTOR Inhibition and Kidney Diseases. Transplantation 102, S32 (2018).

58. McConnachie, D. J., Stow, J. L. & Mallett, A. J. Ciliopathies and the Kidney: A Review. Am. J. Kidney Dis. 77, 410–419 (2021).

59. Singh, P., Harris, P. C., Sas, D. J. & Lieske, J. C. The genetics of kidney stone disease and nephrocalcinosis. Nat. Rev. Nephrol. 18, 224–240 (2022).

60. Weeks, E. M. et al. Leveraging polygenic enrichments of gene features to predict genes underlying complex traits and diseases. 2020.09.08.20190561 Preprint at 10.1101/2020.09.08.20190561 (2020).

61. Li, Y., Cheng, C. N., Verdun, V. A. & Wingert, R. A. Zebrafish nephrogenesis is regulated by interactions between retinoic acid, mecom, and Notch signaling. Dev. Biol. 386, 111–122 (2014).

62. Hoyt, P. R. et al. The Evil proto-oncogene is required at midgestation for neural, heart, and paraxial mesenchyme development. Mech. Dev. 65, 55–70 (1997).

63. Marneros, A. G. AP-2β/KCTD1 Control Distal Nephron Differentiation and Protect against Renal Fibrosis. Dev. Cell 54, 348–366.e5 (2020).

64. Moser, M. et al. Enhanced apoptotic cell death of renal epithelial cells in mice lacking transcription factor AP-2β. Genes Dev. 11, 1938–1948 (1997).

65. Werth, M. et al. Transcription factor TFCP2L1 patterns cells in the mouse kidney collecting ducts. eLife 6, e24265 (2017).

66. Lu, W. et al. NFIA Haploinsufficiency Is Associated with a CNS Malformation Syndrome and Urinary Tract Defects. PLOS Genet. 3, e80 (2007).

67. Li, L. et al. FoxO3 activation in hypoxic tubules prevents chronic kidney disease. J. Clin. Invest. 129, 2374–2389 (2019).

68. Langworthy, M., Zhou, B., de Caestecker, M., Moeckel, G. & Baldwin, H. S. NFATc1 Identifies a Population of Proximal Tubule Cell Progenitors. J. Am. Soc. Nephrol. JASN 20, 311–321 (2009).

69. Kumasaka, N., Knights, A. J. & Gaffney, D. J. Fine-mapping cellular QTLs with RASQUAL and ATAC-seq. Nat. Genet. 48, 206–213 (2016).

70. van de Geijn, B., McVicker, G., Gilad, Y. & Pritchard, J. K. WASP: allele-specific software for robust molecular quantitative trait locus discovery. Nat. Methods 12, 1061–1063 (2015).

71. Yang, T. et al. Expression of peroxisomal proliferator-activated receptors and retinoid X receptors in the kidney. Am. J. Physiol. 277, F966–973 (1999).

72. Cao, X. et al. Chromatin accessibility dynamics dictate renal tubular epithelial cell response to injury. Nat. Commun. 13, 7322 (2022).

73. Gasperini, M. et al. A Genome-wide Framework for Mapping Gene Regulation via Cellular Genetic Screens. Cell 176, 377–390.e19 (2019).

74. Wilson, P. C. et al. Multimodal single cell sequencing implicates chromatin accessibility and genetic background in diabetic kidney disease progression. Nat. Commun. 13, 5253 (2022).

75. Qiu, C. et al. Renal compartment-specific genetic variation analyses identify new pathways in chronic kidney disease. Nat. Med. 24, 1721–1731 (2018).

76. Baigent, C. et al. Impact of diabetes on the effects of sodium glucose co-transporter-2 inhibitors on kidney outcomes: collaborative meta-analysis of large placebo-controlled trials. The Lancet 400, 1788–1801 (2022).

77. Wheeler, D. C. et al. A pre-specified analysis of the DAPA-CKD trial demonstrates the effects of dapagliflozin on major adverse kidney events in patients with IgA nephropathy. Kidney Int. 100, 215–224 (2021).

78. Fulco, C. P. et al. Systematic mapping of functional enhancer–promoter connections with CRISPR interference. Science 354, 769–773 (2016).

79. Mumbach, M. R. et al. Enhancer connectome in primary human cells identifies target genes of disease-associated DNA elements. Nat. Genet. 49, 1602–1612 (2017).

80. Geng, H. et al. Inhibition of Autoregulated TGFβ Signaling Simultaneously Enhances Proliferation and Differentiation of Kidney Epithelium and Promotes Repair Following Renal Ischemia. Am. J. Pathol. 174, 1291–1308 (2009).

81. Park, J. S. et al. N-myc downstream regulated gene 1 (ndrg1) functions as a molecular switch for cellular adaptation to hypoxia. eLife 11, e74031 (2022).

82. Bartsch, D. et al. mRNA translational specialization by RBPMS presets the competence for cardiac commitment in hESCs. Sci. Adv. 9, eade1792 (2023).

83. Identification of RBPMS as a mammalian smooth muscle master splicing regulator via proximity of its gene with super-enhancers | eLife. https://elifesciences.org/articles/46327.

84. Taguchi, K. et al. Cyclin G1 induces maladaptive proximal tubule cell dedifferentiation and renal fibrosis through CDK5 activation. J. Clin. Invest. 132, e158096 (2022).

85. Morris, J. A. et al. Discovery of target genes and pathways at GWAS loci by pooled single-cell CRISPR screens. Science 380, eadh7699 (2023).

86. Schlosser, P. et al. Meta-analyses identify DNA methylation associated with kidney function and damage. Nat. Commun. 12, 7174 (2021).

87. Corces, M. R. et al. An improved ATAC-seq protocol reduces background and enables interrogation of frozen tissues. Nat. Methods 14, 959–962 (2017).

88. Smith, J. P., et al. PEPATAC: an optimized pipeline for ATAC-seq data analysis with serial alignments. NAR Genomics Bioinforma. 3, lqab101 (2021).

89. Heinz, S. et al. Simple combinations of lineage-determining transcription factors prime cis-regulatory elements required for macrophage and B cell identities. Mol. Cell 38, 576–589 (2010).

90. Granja, J. M. et al. ArchR is a scalable software package for integrative single-cell chromatin accessibility analysis. Nat. Genet. 53, 403–411 (2021).

91. Weirauch, M. T. et al. Determination and inference of eukaryotic transcription factor sequence specificity. Cell 158, 1431–1443 (2014).

92. Das, S. et al. Next-generation genotype imputation service and methods. Nat. Genet. 48, 1284–1287 (2016).

93. Fuchsberger, C., Abecasis, G. R. & Hinds, D. A. minimac2: faster genotype imputation. Bioinformatics 31, 782–784 (2015).

94. Taliun, D. et al. Sequencing of 53,831 diverse genomes from the NHLBI TOPMed Program. Nature 590, 290–299 (2021).

95. Fornes, O. et al. JASPAR 2020: update of the open-access database of transcription factor binding profiles. Nucleic Acids Res. 48, D87–D92 (2020).

96. Grant, C. E., Bailey, T. L. & Noble, W. S. FIMO: scanning for occurrences of a given motif. Bioinforma. Oxf. Engl. 27, 1017–1018 (2011).

97. Gazal, S. et al. Functional architecture of low-frequency variants highlights strength of negative selection across coding and non-coding annotations. Nat. Genet. 50, 1600–1607 (2018).

98. Bycroft, C. et al. The UK Biobank resource with deep phenotyping and genomic data. Nature 562, 203–209 (2018).

99. Leeuw, C. A. de, Mooij, J. M., Heskes, T. & Posthuma, D. MAGMA: Generalized Gene-Set Analysis of GWAS Data. PLOS Comput. Biol. 11, e1004219 (2015).

100. Auton, A. et al. A global reference for human genetic variation. Nature 526, 68–74 (2015).

101. Karczewski, K. J. et al. Systematic single-variant and gene-based association testing of thousands of phenotypes in 394,841 UK Biobank exomes. Cell Genomics 2, 100168 (2022).

102. Zou, Y., Carbonetto, P., Wang, G. & Stephens, M. Fine-mapping from summary data with the “Sum of Single Effects” model. PLOS Genet. 18, e1010299 (2022).

103. Kelley, D. R. et al. Sequential regulatory activity prediction across chromosomes with convolutional neural networks. Genome Res. 28, 739–750 (2018).

104. Sanson, K. R. et al. Optimized libraries for CRISPR-Cas9 genetic screens with multiple modalities. Nat. Commun. 9, 5416 (2018).

105. Horlbeck, M. A. et al. Compact and highly active next-generation libraries for CRISPR-mediated gene repression and activation. eLife 5, e19760 (2016).

106. Alerasool, N., Segal, D., Lee, H. & Taipale, M. An efficient KRAB domain for CRISPRi applications in human cells. Nat. Methods 17, 1093–1096 (2020).

107. 10x Genomics. https://support.10xgenomics.com/single-cell-gene-expression/software/downloads/7.0/.

108. Heaton, H. et al. Souporcell: robust clustering of single-cell RNA-seq data by genotype without reference genotypes. Nat. Methods 17, 615–620 (2020).

109. Hao, Y. et al. Integrated analysis of multimodal single-cell data. Cell 184, 3573–3587.e29 (2021).

110. Replogle, J. M. et al. Combinatorial single-cell CRISPR screens by direct guide RNA capture and targeted sequencing. Nat. Biotechnol. 38, 954–961 (2020).

111. Joseph Replogle. https://github.com/josephreplogle/guide_calling

